# Evaluating and integrating spatial capture-recapture models with data of variable individual identifiability

**DOI:** 10.1101/2020.03.27.010850

**Authors:** Joel S. Ruprecht, Charlotte E. Eriksson, Tavis D. Forrester, Darren A. Clark, Michael J. Wisdom, Mary M. Rowland, Bruce K. Johnson, Taal Levi

## Abstract

Spatial capture-recapture (SCR) models have become the preferred tool for estimating densities of carnivores. Within this family of models are variants requiring identification of all individuals in each encounter (SCR), a subset of individuals only (generalized spatial mark-resight, gSMR), or no individual identification (spatial count or spatial presence-absence). Although each technique has been shown through simulation to yield unbiased results, the consistency and relative precision of estimates across methods in real-world settings are seldom considered. We tested a suite of models ranging from those only requiring detections of unmarked individuals to others that integrate remote camera, physical capture, genetic, and global positioning system (GPS) data into a ‘hybrid’ model, to estimate population densities of black bears, bobcats, cougars, and coyotes. For each species we genotyped fecal DNA collected with detection dogs during a 20-day period. A subset of individuals from each species was affixed with GPS collars bearing unique markings and resighted by remote cameras over 140 days contemporaneous with scat collection. Camera-based gSMR models produced density estimates that differed by less than 10% from genetic SCR for bears, cougars, and coyotes once important sources of variation (sex or behavioral status) were controlled for. For bobcats, SCR estimates were 33% higher than gSMR. The cause of the discrepancies in estimates was likely attributable to challenges designing a study compatible for species with disparate home range sizes and the difficulty of collecting sufficient data in a timeframe in which demographic closure could be assumed. Unmarked models estimated densities that varied greatly from SCR, but estimates became more consistent in models wherein more individuals were identifiable. Hybrid models containing all data sources exhibited the most precise estimates for all species. For studies in which only sparse data can be obtained and the strictest model assumptions are unlikely to be met, we suggest researchers use caution making inference from models lacking individual identity. For best results, we further recommend the use of methods requiring at least a subset of the population is marked and that multiple datasets are incorporated when possible.

## Introduction

Population abundance is a state variable of paramount importance in both applied and basic ecology but remains a challenge to estimate for most free-ranging animal populations. This is particularly the case for terrestrial carnivore species that are cryptic and occur at low densities (Wilson and Delahay 2001). Many statistical models accounting for imperfect detection have been used to estimate abundance, but the data required for each method vary along a continuum of identifiability of individual animals, ranging from no individual recognition to full individual recognition. For situations in which no animals can be uniquely identified, models to estimate abundance have been developed based on presence-absence of unmarked individuals in a sampling session (Royle and Nichols 2003, Ramsey et al. 2015) or counts of unmarked individuals (Buckland et al. 1993, Royle 2004, Rowcliffe et al. 2008, Chandler and Royle 2013, Moeller et al. 2018). Other models rely on partially-observable encounter histories when a subset of the population is identifiable (e.g. White 1996, McClintock and White 2009, Sollmann et al. 2013a, Augustine et al. 2018), and the most data-intensive models utilize fully-observable encounter histories such that all individuals can be uniquely identified (e.g., Otis et al. 1978, Nichols 1992, Efford 2004, Royle et al. 2013). Collecting higher resolution data where individuals are identifiable is substantially more challenging and costly, but models based on these data may produce more robust estimates of abundance particularly in situations with suboptimal study designs or when model assumptions cannot strictly be upheld (Chandler and Royle 2013, Augustine et al. 2019). Knowing what type and how much data to collect to achieve satisfactory results is therefore a central challenge to estimate population abundance for conservation, management, or research objectives.

Spatial capture-recapture (SCR; Efford 2004, Borchers and Efford 2008, Royle and Young 2008, Royle et al. 2013) has emerged as the most prominent method of abundance estimation for carnivores over the past five years (Appendix S1: Fig. S1). SCR models exploit the spatial locations of animal detections and assume the rate or probability an individual is detected is highest at its home range center and declines as distance from that location increases (Royle and Young 2008). The spatially-referenced detections of an individual allow the estimation of the animal’s activity center, and the number of estimated activity centers yields an estimate of abundance. Population density, often preferable to abundance, is calculated by dividing the estimated number of activity centers by the area of the state space which eliminates the arbitrarily-defined effective sampling area required in traditional capture-mark-recapture approaches (Royle et al. 2013). Furthermore, SCR can accommodate individual and trap-level covariates to explain spatial and individual variation in detection rates.

SCR models are well-suited for noninvasive genetic sampling in which individuals are uniquely identified, typically using DNA from scat or hair samples, to produce fully-observable encounter histories (Gardner et al. 2009, Kery et al. 2011, Morin et al. 2016, 2018). Scat-detection dogs in particular have become a common tool to efficiently collect scat on large landscapes within a narrow time window to ensure demographic closure even for species that occur at low densities (Wasser et al. 2004, 2011). While effective for SCR density estimation, scat detection requires highly-trained dogs and specialization in genetic techniques or the resources to contract out those tasks. As a more inexpensive and accessible alternative, wildlife researchers increasingly use remote cameras to study and monitor wildlife. However, cameras can only be used for conventional SCR analyses when all individuals of a species possess natural markings (e.g. unique pelage patterns) that permit individual identification (an exception is the spatial partial identity model; Augustine et al. 2018).

Spatial mark-resight (SMR) and generalized spatial mark-resight (gSMR) models are variants of SCR that allow spatially-explicit density estimation when only a subset of the population can be uniquely identified (Chandler and Royle 2013, Sollmann et al. 2013a, Whittington et al. 2018). A common example is identification of individual study animals that are marked with GPS (global positioning system) collars or other telemetry devices. For many species it would be inefficient to capture and mark animals solely for the purpose of density estimation; however, many researchers and wildlife management agencies routinely monitor populations with telemetry devices. In these cases, resighting marked animals on an array of remote cameras may offer a relatively straightforward option for estimating density at little extra cost (Whittington et al. 2018).

Due to the challenge of individually marking animals, spatial presence-absence (SPA; Ramsey et al. 2015) and spatial count (SC) or ‘unmarked SCR’ models (Chandler and Royle 2013) have been developed to exploit spatial autocorrelation in counts of unmarked animals to make inference on their activity centers, and by extension, population density. The attractiveness of these approaches is that they require no individual identification so the population need not be artificially nor naturally marked. Theoretically, the only data required for SPA or SC models are spatially-referenced binary detections or counts of animal detections, respectively, which allows use of standard camera trapping datasets that can be collected noninvasively without the cost and specialized equipment needed for genotyping. Although Chandler and Royle (2013) caution researchers to mark a subset of the population whenever possible, others have successfully estimated population densities using datasets lacking any individual recognition (e.g., Evans and Rittenhouse 2018). If this were universally true, SC methods would represent a major advance in our ability to rapidly and affordably count carnivores.

These three model types (i.e., SC, SMR/gSMR, and SCR) require data of vastly different levels of individual identifiability which raises concerns over the consistency of results one may observe if using multiple methods. Understanding the relative performance of these commonly-used density estimators is critical to guide future population monitoring for conservation and management. We hypothesized, *a priori*, that genetic SCR methods would yield the most robust density estimates because they require fully-observable encounter histories. As the use of gSMR, SMR, and SC methods proliferates in the literature, however, it is important to evaluate their credibility with respect to the ‘gold standard’ SCR approach. A variety of additional data sources can be incorporated into these models, which could potentially increase the concordance of results from unmarked models to fully-marked SCR models while also increasing precision and accuracy. For example, all of these models can be informed by telemetry data in various capacities (Sollmann et al. 2013a) to provide independent information on detection parameters, and the SMR model should additionally be informed by the capture process used to mark animals to reduce bias (i.e. generalized SMR; Whittington et al. 2018). Since research projects and monitoring programs have limited budgets, it is also critical to know how much and what type of data must be collected to achieve satisfactory results.

Here we use a unique dataset consisting of 1) genotyped scats located by detection dogs, 2) images of both marked and unmarked animals from an array of remote cameras, 3) encounter histories from live-trapping, and 4) GPS collar locations to estimate the abundances of four carnivore species: black bears (*Ursus americanus*), bobcats (*Lynx rufus*), cougars (*Puma concolor*), and coyotes (*Canis latrans)*. Our objective is to evaluate a suite of existing spatial density models across a range of individual identifiability to evaluate agreement in results and precision under realistic data collection scenarios. In addition, we present a new hybrid model that can incorporate SCR and gSMR data sources (i.e. including genotypes, the physical marking process, camera data, and GPS data) into a single hierarchical model, and assess the resulting gains in precision. Notably our model comparisons occur in the context of a real system rather than simulation. Each species varies in its life history, from highly territorial but group living (coyotes), to non-territorial solitary individuals (black bears), to solitary territorial individuals with territory size varying by sex (bobcat females are territorial while males are not; cougar males are territorial while females are not). By evaluating a suite of methods with different data requirements, our findings directly address how population densities of terrestrial carnivores can most efficiently be estimated, and whether the investment in genotyping or marking and resighting animals is warranted given the availability of models that do not rely on individual identification.

## Methods

### Study Area

Our study was conducted in and adjacent to the Starkey Experimental Forest and Range (hereafter Starkey) in the Blue Mountains of northeastern Oregon (Fig. 1). Starkey is bounded by a 2.4-meter fence, enclosing 100.7 km^2^, which prevents the passage of large herbivores (Rowland et al. 1997); however, carnivores are undeterred by the fence and regularly cross in and out of the enclosure (this study, Oregon Department of Fish and Wildlife, unpublished data). Starkey is part of the Wallowa-Whitman National Forest and is administered by the US Forest Service. Land immediately adjacent to Starkey is predominantly public and managed by the US Forest Service (Wallowa-Whitman and Umatilla National Forests) but also includes private inholdings. The study area contains a mosaic of Ponderosa pine (*Pinus ponderosa*) and mixed pine-fir forests (*Pinus*, *Abies* and *Pseudotsuga* spp.), interspersed with grasslands dominated by native bunchgrasses (*Poa*, *Danthonia*, and *Pseudoroegneria* spp.) and invasive annual grasses (*Bromus* and *Ventenata* spp.) (Rowland et al. 1997). Elevation ranges between 1,122 and 1,500 m within Starkey (Rowland et al. 1997). See Wisdom et al. (2005) for additional details about Starkey.

**Fig. 1:**
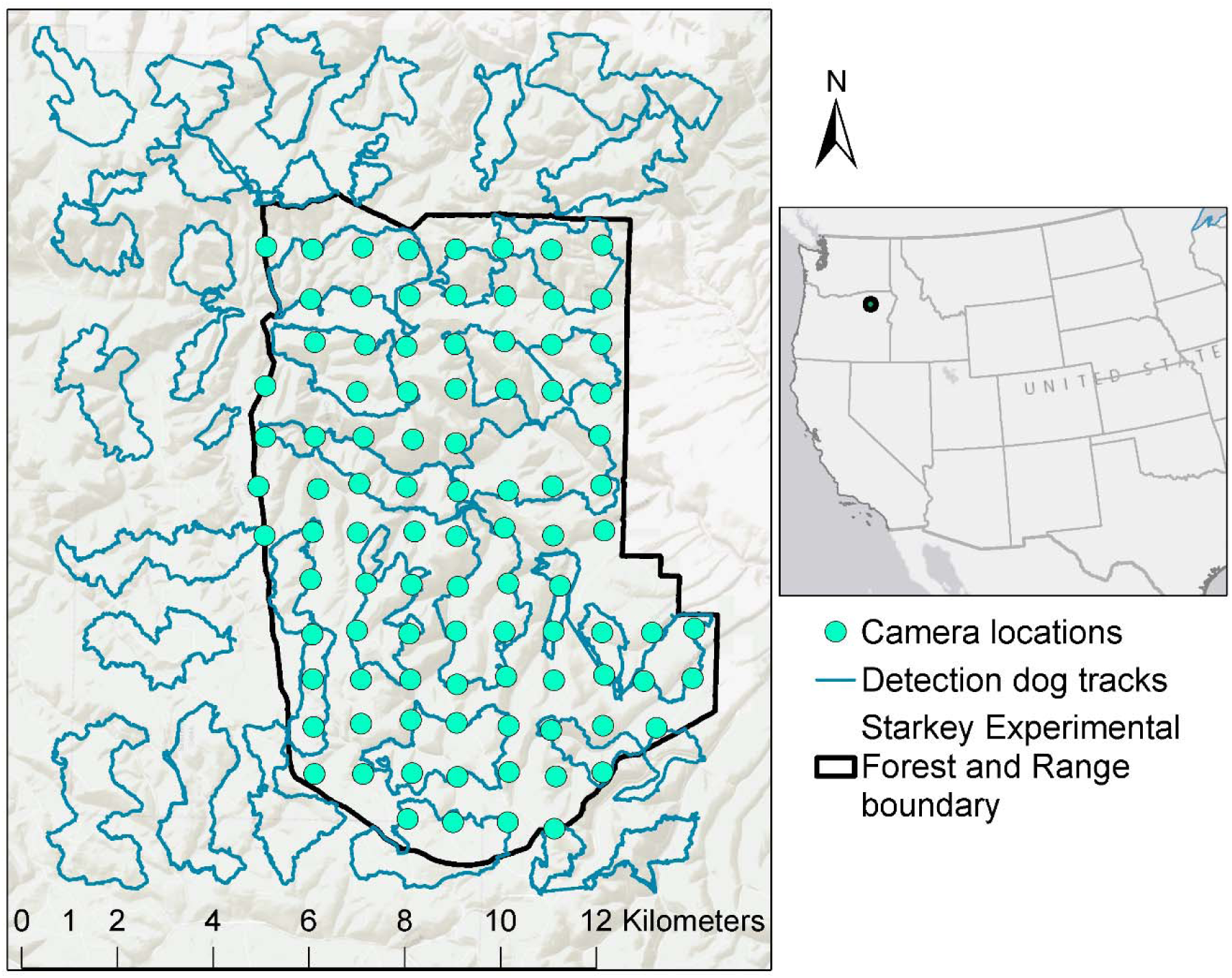
The study area in northeast Oregon in which the densities of black bears, bobcats, cougars, and coyotes were estimated using a suite of spatially-explicit models fit to data collected by remote cameras and noninvasive genetic sampling. The black polygon represents the Starkey Experimental Forest and Range boundary, green points are camera locations, and blue lines are scat detection dog survey tracks.

### Data collection

#### Scat dataset

Three teams of scat detection dogs from the University of Washington Conservation Canine program surveyed a 224 km^2^ study area encompassing all of Starkey and roughly an equal area immediately to the north and west of the enclosure between 6 and 26 June 2017 (Fig. 1). The area surveyed was composed of 56 grid cells each with an area of 4 km^2^. Detection dogs surveyed 6–8 km linear distance within each cell to distribute effort across the study area. Dog handlers were not given specific survey routes but were encouraged to follow natural travel corridors such as ridgelines, saddles, drainage bottoms, game trails, and fencelines. No more than 50% of the distance traveled per grid cell was permitted to be on linear features such as trails or roads. When scats were located, the handler recorded the GPS position and placed the entire scat in triplicate paper bags (Fig. 2a). Within 72 hours of collection, scats were desiccated in a drying oven for 24 hours at 40°C (Murphy et al. 2000). Detection dogs in our study were trained to locate black bear, coyote, cougar, and bobcat scats.

**Fig. 2:**
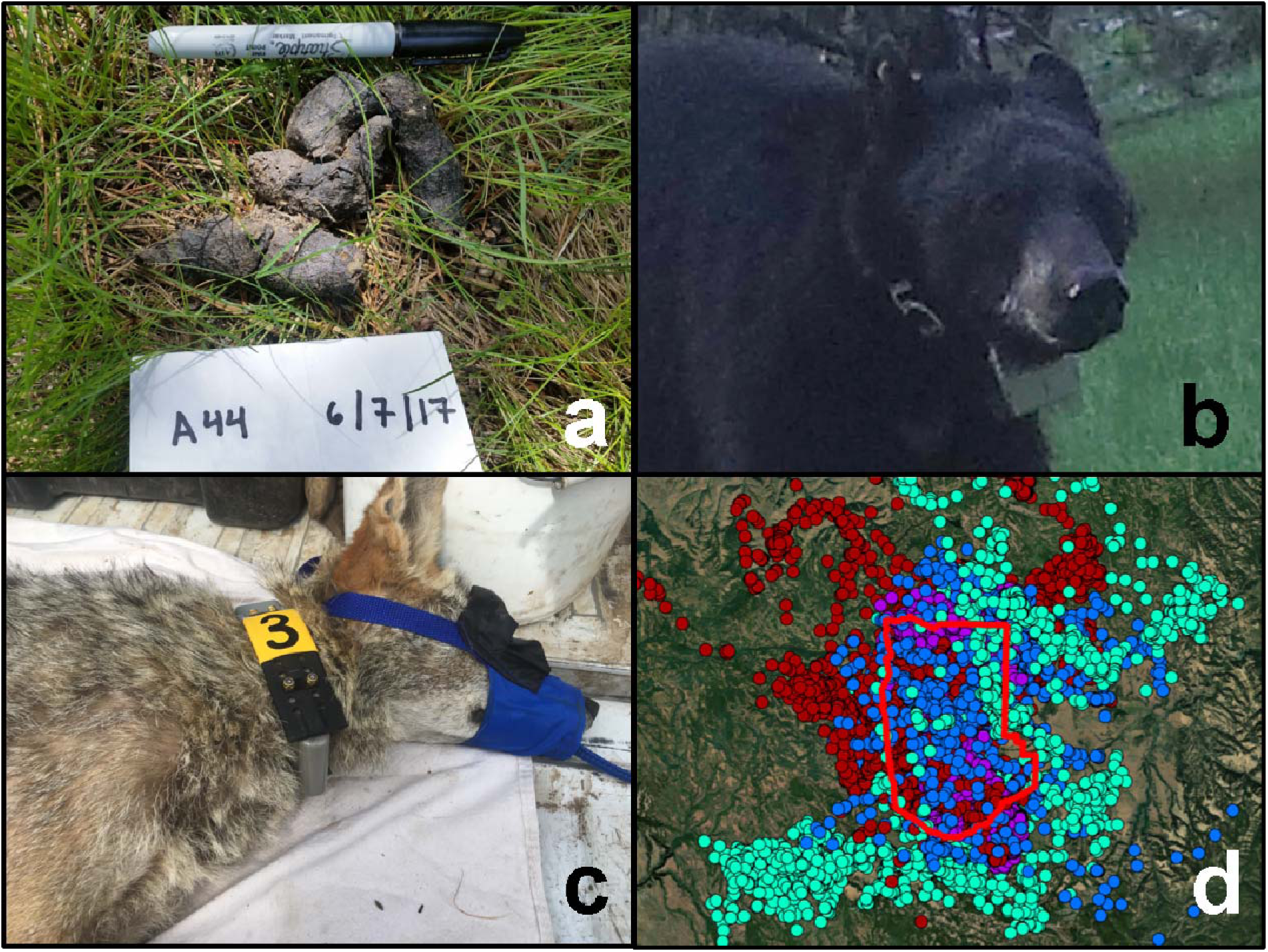
a) Scat located by detection dogs subsequently analyzed for species ID and genotyped to determine individual identity. b) A GPS-collared bear detected on remote camera. The number on the collar belting and location of the ear tag permitted individual identification of this animal. c) Immobilized coyote fit with GPS collar. Note the colored shrinkwrap around collar banding and unique number aiding in individual identification when photographed on remote cameras. d) GPS collar positions for coyotes (light green), cougars (blue), bobcats (purple), and bears (red) during the time sampling was conducted. The red outline is the Starkey Experimental Forest and Range boundary.

We used genotyping by amplicon sequencing (Eriksson et al. 2020) to identify unique individuals with single-nucleotide polymorphisms (SNPs), a process that uses the power of high-throughput sequencers to yield high genotype success rates and low error rates. Coyote scats were genotyped following methods in Eriksson et al. (2020). We genotyped bobcat, cougar, and black bears scat using the same approach but with novel primers designed and tested specifically for this study (Appendix S2). Prior to genotyping, the identity of the defecator for each scat was identified using DNA metabarcoding of ∼100 base pairs of the mitochondrial 12S gene region as part of a separate diet study (primers used in Eriksson et al. 2020, Massey et al. 2020, and modified from Riaz et al. 2011). See Appendix S2 for SNP discovery methods and Eriksson et al. (2020) for detailed primer design and genotyping protocols.

#### Camera dataset

A 1- x 1-km grid of 94 unbaited remote cameras (Bushnell TrophyCam Aggressor, Overland, KS) was placed inside the Starkey fence perimeter and was operational between April and September 2017 (Fig. 1). All cameras were pointed north. We visited each camera station every 6-8 weeks to change batteries and download data. Each camera was set to take a burst of 3 photos when triggered.

Photos were tagged to species or to individual when animals were uniquely identifiable due to GPS collars and ear tags. We used the open source photo management software DigiKam (www.digikam.org) for tagging and database management and the R library camtrapR (Niedballa et al. 2016) to extract metadata from photos. We censored photos in which we could not determine whether an animal was wearing a GPS collar or if we could not verify the identity of an animal photographed with a GPS collar. However, these records constituted < 5% of all detections. Because animals were typically photographed more than once during a visit, we recorded information for each independent photo sequence and considered photo sequences > 30 minutes from the next detection of the same species to be independent.

#### Telemetry dataset

We captured and GPS-collared a sample of individuals from each of the four carnivore species as part of concurrent research on predator-prey relationships. Coyotes, bobcats, and bears were captured inside Starkey only, and cougars were captured in Starkey plus a buffer of approximately 20 km which was encompassed by the state space. Coyotes were captured using padded foothold traps (Oneida Victor SoftCatch No. 3, Euclid, OH) with a tranquilizer tab (Balser 1965) containing 50 mg propriopromazine hydrochloride attached to each trap to subdue captured animals until researchers arrived. Captured animals were immobilized with tiletamine-zolazepam (Telazol®) at a concentration of 10 mg/kg. A GPS collar (Lotek MiniTrack, Lotek Wireless Inc., Newmarket, ON, Canada or Vectronic Vertex, Vectronic Aerospace GmbH, Berlin, Germany) was placed on each adult coyote and was programmed to record locations every 2 or 3 hours.

We captured bobcats using cage traps baited with visual and olfactory attractants. Captured bobcats were administered ketamine (10 mg/kg) and xylazine (1.5 mg/kg) for immobilization, and upon release, yohimbine (0.125 mg/kg; Yobine®) was given as an antagonist for xylazine. Each bobcat was fit with a GPS collar (Lotek MiniTrack, Lotek Wireless Inc., Newmarket, ON, Canada) scheduled to take fixes every 2 hours.

Black bears were captured using culvert traps or padded foot snares (Lemieux and Czetwertynski 2006). Captured bears were immobilized with Telazol® at a concentration of 7 mg/kg. Bears were fit with GPS collars (Lotek GPS 7000 or LiteTrack Iridium 420, Lotek Wireless Inc., Newmarket, ON, Canada) that recorded positions every 15 min to 2 hours.

We captured cougars using trained pursuit hounds. We systematically searched roads for fresh cougar tracks in snow and when located, released hounds to pursue tracks until the cougar was treed. Dogs were allowed to pursue tracks of any individual cougar. When treed, cougars were immobilized with ketamine (10 mg/kg) and xylazine (2 mg/kg) via dart gun, and before release were administered yohimbine (0.125 mg/kg; Yobine®) as an antagonist for xylazine. A GPS collar (Lotek GPS 4400S, IridiumTrack M, Lotek Wireless Inc., Newmarket, ON, Canada or Vectronic Vertex Lite, Vectronic Aerospace GmbH, Berlin, Germany) was placed on each cougar and was programmed to record locations every 3 hours.

We permanently marked the belting of GPS collars with unique numbers used to identify individuals on camera (Fig. 2b). When possible (i.e., for the larger collars), we also affixed sections of colored heat-shrink plastic tubing over the collar belting to assist with identification (Fig. 2c). We ear-tagged each captured animal such that females received ear tags on the left ear and males on the right ear which provided an additional cue when identifying GPS-collared animals on camera.

We collected biological samples from each captured individual for SNP primer development and to match the genotypes of GPS-collared animals with genotypes detected by scat detection dogs. For each animal we sampled some combination of tissue (ear-tag punch), hair, and scat (see Eriksson et al. 2020 for additional details). All animal handling was performed in accordance with protocols approved by the USDA Forest Service, Starkey Experimental Forest Institutional Animal Care and Use Committee (IACUC No. 92-F-0004; protocol #STKY-16-01) and followed the guidelines of the American Society of Mammalogists for the use of wild mammals in research (Sikes 2016).

### Modelling Approach

Our approach was to fit a suite of spatially-explicit models across a spectrum of data resolutions (Table 1) for each species to evaluate the value of each data source and to test whether incorporating multiple data sets improved precision. Each model considered here is a variation of a conventional SCR model (Royle and Young 2008) in which the detection probability *p_i,j_* (or rate *λ_i,j_*) decays as a function of the Euclidean distance between the detector *j* and the activity center *s* for individual *i*, *d_i,j_*, to yield

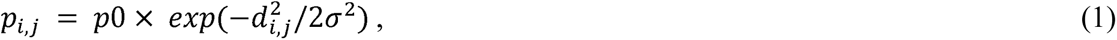

**Table 1:**
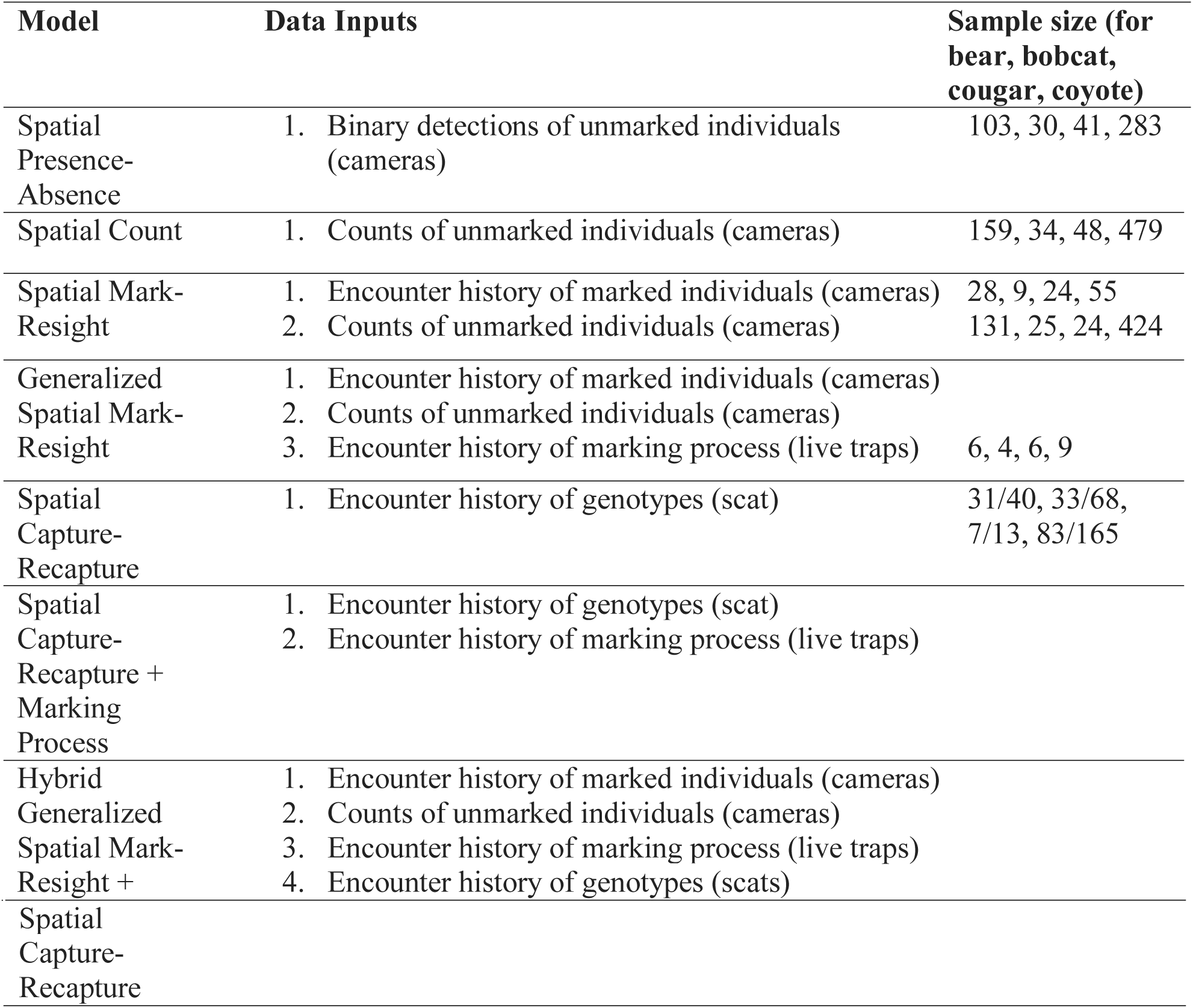
Description of data requirements for each model evaluated to estimate densities of four carnivore species in northeast Oregon. The data collection sources used in this study are given in parentheses. Models are ordered in terms of increasing data requirements. Sample sizes for the number of independent detections for each data collection method are given in the last column, separated by commas, for black bears, bobcats, cougars, and coyotes, respectively. For sample sizes of genetic samples, we provide both the number of unique individuals and number of total samples, respectively, separated by a slash (e.g. 83/165 indicates 83 unique coyotes were detected from 165 genotyped scats). For brevity we only present sample sizes the first time a certain data type is introduced.

where *p*0 is the baseline detection probability when the detector is located exactly at the activity center of the animal, and is a spatial scale parameter related to home range size that determines the rate at which detection probability declines with distance from the activity center. The location of each activity center (*s_i_*) is latent but estimated from the locations in which detections and non-detections occurred.

The observed encounter frequencies *y_i,j_* can be modeled as either a binomial or Poisson random variable, depending on whether the detection process produces binary or count outcomes,

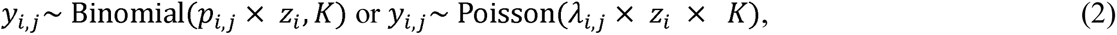

where *K* is the number of sampling occasions, and *z_i_* is the data augmentation parameter (Royle et al. 2007) that takes the values of 0 or 1 for *i* = 1, 2…*M*, where *M* is the number of known individuals plus some number of all zero encounter frequencies representing individuals in the population not detected during sampling. The value of *M* must be chosen to be larger than the maximum possible population size and is sufficiently large when it does not affect the estimation of *N*. Data augmentation is used to estimate the number of animals in the state space, *S* (an area prescribed to be sufficiently large such that animals with activity centers at the outermost regions of this area have a negligible probability of being detected). *z_i_* determines whether each hypothetical individual in the augmented population is real or not and is governed by a Bernoulli distribution,

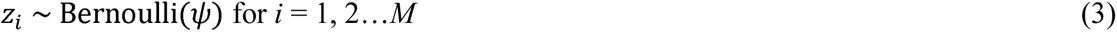

in which ψ is the expected number of activity centers divided by *M*. The estimated population size *N* (i.e. the number of activity centers in the state space) is then determined by summing over all *z_i_*,

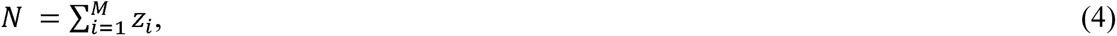

and density *D* is obtained by dividing the population size by the area of the state space, *A*,

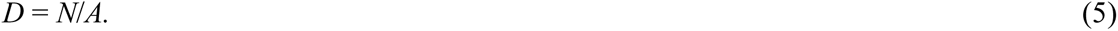

To ensure that the specification of the state space and number of augmented individuals is sufficiently large, *M* must be substantially higher than the estimated *N*, the upper bound of ψ must not be near 1, and the buffer around the survey area forming the state space must be > 2.5 times σ (Royle et al. 2013).

We assumed a homogenous point process for the distribution of activity centers (*s_i_*) such that the *x* and *y* dimensions of each activity center were uniformly distributed within the defined limits of the state space:

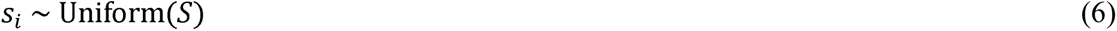

Each model subsequently described is based on the same framework as the conventional SCR model described above but incorporates variations due to different data sources or resolutions which are described next.

#### Spatial Count or ‘unmarked SCR’

The spatial count model (Chandler and Royle 2013) does not require individual recognition, so the encounter frequency *y_i,j_* is replaced by a vector of counts of unmarked individuals at each detector (*n_j_*) and encounter rate *λ* replaces *p*. The counts are now modeled as a Poisson random variable,

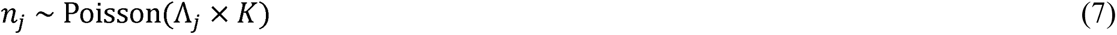

where

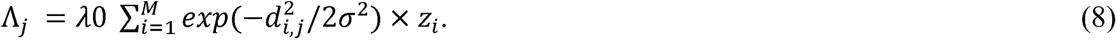

#### Spatial Presence Absence

The Poisson distribution assumes that the mean is equal to the variance, but when animal detections are aggregated in time and/or space the variance may greatly exceed the mean. This overdispersion could bias SC results due to the violation of the model assumption. The spatial presence-absence model (SPA; Ramsey et al. 2015) is similar to the spatial count model, but only detection-non detection data are utilized so a binomial distribution replaces the Poisson. By reducing counts to binary detections and using a binomial distribution, the SPA model may be more appropriate in situations with overdispersion.

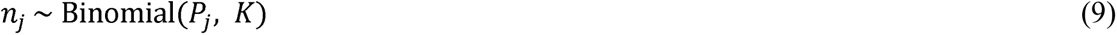

where *n_j_* ∈ {0,1} and

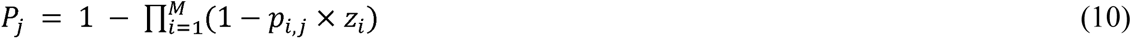

Above, *P_j_* is the probability that at least one individual is detected in detector *J*, and *p_i,j_* is the detection probability for individual *i* at detector *J* which like the other models decreases as a function of the distance between the activity center and the trap according to a half-normal function (as in equation 1).

#### Spatial Mark-Resight

Spatial mark-resight models (Chandler and Royle 2013, Sollmann et al. 2013a) combine SCR and SC. Here, a subset of the population is individually recognizable due to either natural or artificial markings but the remaining individuals are observed without individual identity. The Poisson-distributed encounter history:

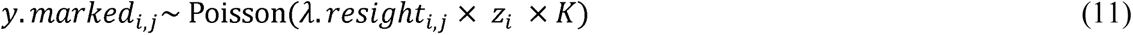

is obtained for marked individuals and the resulting expected resighting rate:

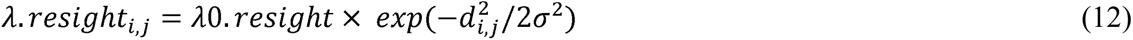

is applied to the counts of unmarked individuals *n_j_* by replacing *λ*0 in Eq. 8 with *λ*0*resight* ’. Assumptions of this model are that marked and unmarked individuals have the same rate of detection and are always accurately identified as marked or not.

#### Generalized Spatial Mark-Resight

Generalized spatial mark-resight (Whittington et al. 2018) is identical to the conventional SMR described above but incorporates a submodel for the marking (i.e. physical capture and tagging) process and is therefore suitable only when a portion of the population is captured and marked. Generalized SMR alleviates bias introduced due to heterogeneity of individual encounter rate arising from situations when the marked portion of the population resides nearer to the resighting detectors (and are therefore detected more frequently) than unmarked individuals at the periphery of the state space (Whittington et al. 2018). That is, it accounts for the fact that the marked segment of the population is seldom distributed randomly throughout the state space (Whittington et al. 2018). The marking process submodel is itself a SCR model for the encounter histories of individuals captured and marked:

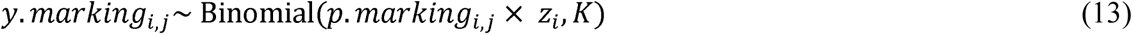

where the probability of capturing individual *i* in trap *j* similarly decays with distance according to

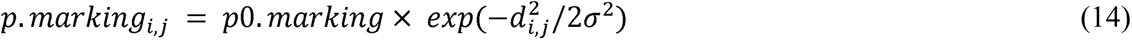

The animal capture and marking process and the resighting of marked animals on cameras shares information on *z_i_*, *s_i_*, and possibly σ, which has the potential to increase model performance and precision. Generalized spatial mark-resight should be used whenever possible because conventional SMR models have been shown to be biased (Whittington et al. 2018).

#### Hybrid Generalized Spatial Mark-Resight + Spatial Capture-Recapture

When genetic SCR data are obtained concurrently with a capture and marking effort, and the marked individuals are resighted on an array of detectors (e.g. trail cameras), we propose data from all three processes can be modeled together in a ‘hybrid’ gSMR+SCR model (Fig. 3). This model incorporates equations 1-6, and 11-14. This data-intensive model thus utilizes multiple encounter methods (physical capture, genetics, and cameras) and requires that the identities of captured and marked individuals can always be reconciled when encountered in the genetic dataset, so captured animals must always be genotyped. Further, the encounter histories from each of the three data sources must be aligned such that each individual known from either the marking/resighting process or scat detection process corresponds to the same individual, *i* in the combined model. However, not all known individuals are likely to be detected in all data sources, so all zero detection histories must be given to known individuals when not encountered in a certain encounter method. For marked animals which are known to exist because they were physically captured, the possible outcomes of the detection process are 1) detected by neither camera nor genetics, 2) detected by camera only, 3) detected by genetics only, or 4) detected by both camera and genetics. For unmarked animals (i.e. those not physically captured), the possible outcomes of the detection process are 1) detected by neither camera nor genetics, 2) detected by genetics only (and without a genotype matching any of the marked individuals), 3) detected by camera only without individual identity, and 4) detected by camera and genetics (and without the possibility of deterministically linking the genotype to the photographed individual). Information on *z_i_*, *s_i_*, and σ can be shared, and baseline detection rates can be informed by the different data sources when an individual is detected by one but not all detection methods. For example, knowing a captured animal exists but was not detected by genetics can help inform the probability of detection in the genetic SCR component. Similarly, knowing a captured animal exists but was not photographed informs the detection probability for the SMR model component.

**Fig. 3:**
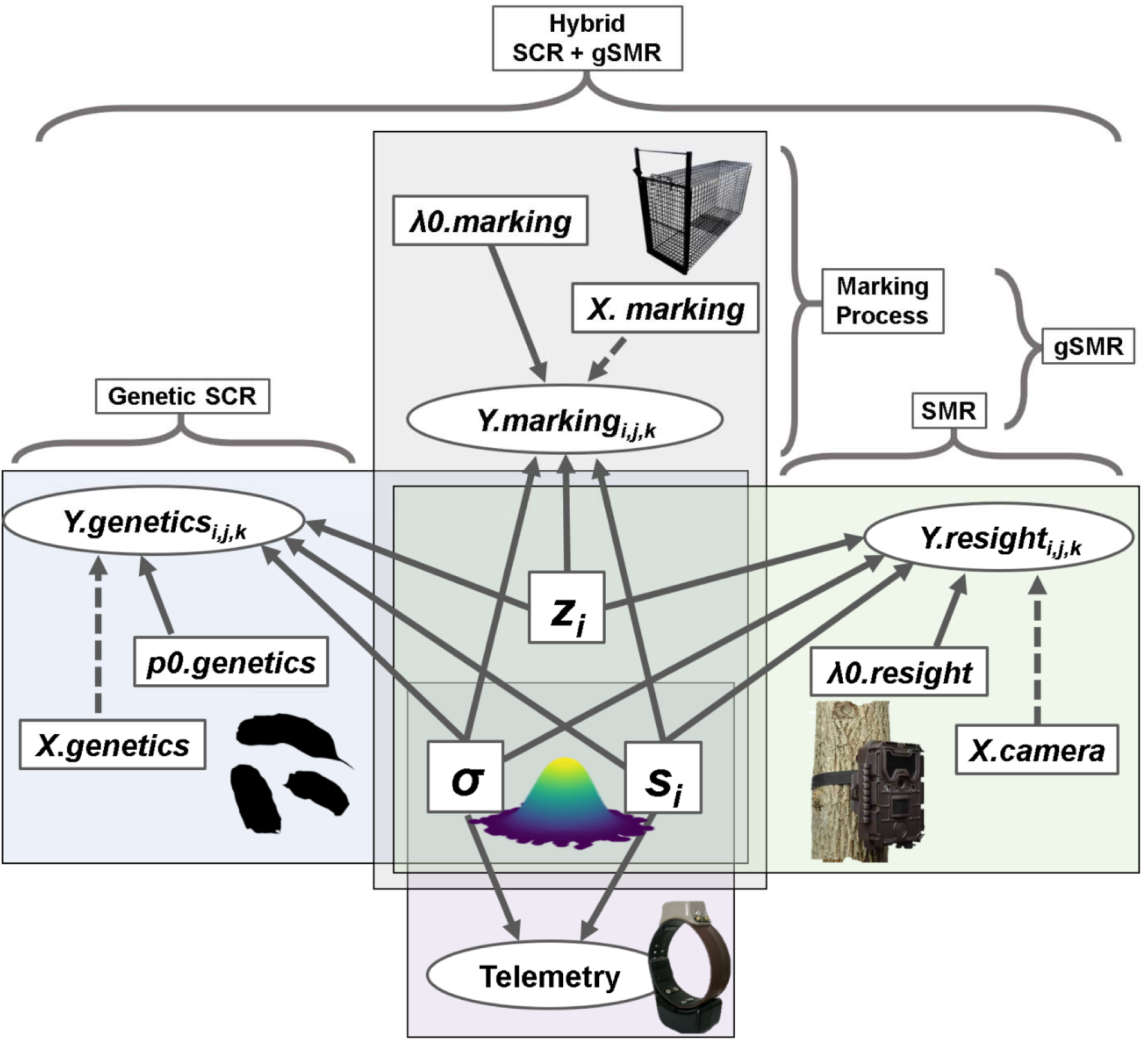
A directed acyclic graph (DAG) displaying the inputs (labeled outside curly brackets) and relationships between data and parameters for the hybrid gSMR+SCR (generalized spatial mark-resight and spatial capture-recapture) model. Dependent variables (i.e. detection data) are represented by ovals. Solid lines connect parameters to data and independent variables are connected to dependent variables by dotted lines. Parameters appearing inside overlapping boxes are jointly estimated across different data sources and submodels. Telemetry data can optionally be incorporated into every model. Note that gSMR is composed of 1) conventional spatial mark-resight (SMR) and 2) the marking process, a SCR model describing the process by which a subset of animals are physically captured and marked allowing them to be resighted.

The hybrid gSMR+SCR model, below with the same distributional assumptions and priors as in our empirical study, has the following joint posterior distribution:

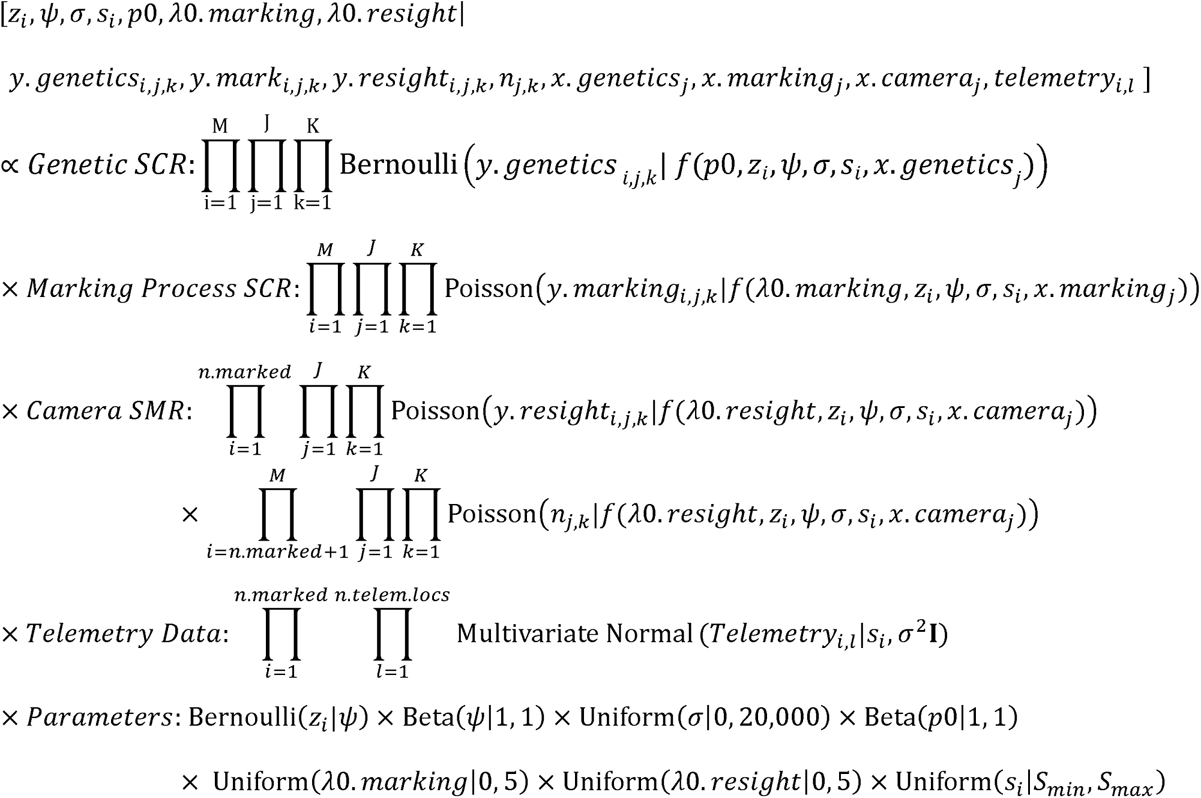

where *x* corresponds to the geographic coordinates of the detectors of a given data source. A directed acyclic graph (DAG) of the hybrid gSMR+SCR model is presented in Fig. 3.

#### Incorporating telemetry data

If some animals within the population are fit with GPS collars (or other telemetry methods) during the time sampling is conducted, the telemetry positions can be incorporated into any of the models previously described (Royle et al. 2013, Sollmann et al. 2013a). This is achieved by modelling the *l* GPS positions as a two-dimensional multivariate normal distribution (i.e. bivariate normal) centered on the activity center coordinates of marked individual *i* with no covariance (Garton et al. 2001):

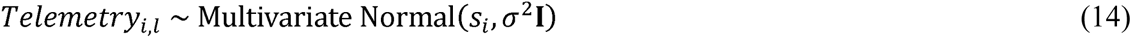

Thus, the incorporation of telemetry data informs the location of the activity center *s* for telemetered individuals, the scale parameter in the detection function, and baseline detection rate or probability. However, in the spatial count models, individual recognition is not known, which prevents the GPS data from being deterministically linked to specific individuals, so in those models the telemetry data typically only inform σ.

#### Application

For each of the four study species (black bear, bobcat, cougar, coyote), we first fit “base” models that did not consider differences in sex or behavioral status in detection parameters. SC models were fit using only camera data. Individual identity was ignored in SC models even when it was known so that we could evaluate an approach that required no individual recognition. Although this was an inefficient use of the information available, our objective was to determine whether investing in the marking and resighting of individuals was necessary. SMR and gSMR models were constructed from the same camera data as the SC models but GPS-collared animals were identified to individual, with the latter method also including the encounter histories from the marking process. SCR models were constructed only from genetic samples (scats) because we could not identify individuals from cameras unless they were artificially marked. The gSMR + SCR hybrid model that we developed combined data from cameras, physical captures to mark animals, and genetic samples. We assumed that camera and scat data were independent because 1) both data collection methods passively sampled the population in different ways and at different sites within the superpopulation, 2) the detection of a scat did not influence the probability of that individual being detected by camera (and vice-versa), and 3) neither camera placements nor scat dog survey routes were influenced by the knowledge gained from the other method.

For every modeling method implemented (Table 1), we fit models both with and without GPS data. When telemetry data were included, we randomly selected 100 GPS locations (Fig. 2d) from each individual to alleviate lack of independence arising from temporal autocorrelation. Because GPS data were collected contemporaneously with camera and genetic sampling, they could be deterministically linked to the detection data of that individual which helped us estimate the activity center for that individual, as well as σ and *p*0 or λ0. The easting and northing locations were modeled as a bivariate normal distribution with no covariance, with a mean of that individual’s activity center (*s*_i_) and standard deviation equal to the detection scale parameter σ.

For the genetic SCR analyses, we used a single occasion because detection dogs only surveyed each grid cell once. We overlaid a 1 x 1 km grid over the survey area and attributed the location of any scat inside that cell to the grid cell center. To control for variation in survey effort, we included the distance walked by scat dog handlers within each grid cell as a covariate influencing *p*0 using a linear model with a logit link. Samples belonging to the same individual that were found in the same grid cell were recorded as a single detection of individual *i*. Because some carnivore species defecate nonrandomly as a means of scent marking, we believe removing recaptures within the same grid cell helped ensure independence of samples.

For camera analyses, we parsed the encounter data into ten 14-day occasions beginning 15 April. We used a trap operation matrix in our detection function to account for varying camera operability so that the probability of detection was zero if a camera was not functional. Similarly, if an animal marked at some point during the sampling period was not available to be detected during one or more occasions due to death or because it was not yet collared, we did not allow that individual to contribute to the likelihood for that occasion.

To assess and compare dispersion across model types, we calculated the coefficient of variation as the posterior standard deviation divided by the posterior mean. We refer to precision as the inverse of the coefficient of variation rather than the inverse of the variance.

#### Model implementation

For models lacking individual identity of all individuals (i.e. all except SCR) we used the marginal model implementation described by Chandler and Royle (2013). We used Markov Chain Monte Carlo (MCMC) to draw samples from the posterior distributions for the parameters of interest. We used JAGS (Plummer 2003) in program R version 3.6.1 (R Development Core Team 2019) and the jagsUI package (Kellner 2015) and NIMBLE (de Valpine et al. 2017, NIMBLE Development Team 2019). For each model we ran 3 chains consisting of 50,000 iterations per chain and discarded the first 15,000 as burn in. We assessed model convergence by visually inspecting traceplots and ensuring the ^-^ values were less than 1.1 (Gelman 1996). If models failed to converge after the first run, we updated them until convergence was satisfactory. For all parameters, we calculated the 95% highest posterior density intervals (HPDI) and 95% Bayesian credible intervals (BCI) (Chen and Shao 1999). For estimates of population density, which may be sensitive to the amount of data augmentation prescribed, we report the posterior mode in the text but also present the posterior means and medians in tables.

## Results

Our capture efforts yielded GPS collars placed on 6 black bears, 4 bobcats, 6 cougars, and 9 coyotes that could be resighted. During the time of camera sampling, 1 cougar died but the remainder of marked individuals remained alive and in the sampling area. However, 3 coyotes were transient, meaning they roamed widely across an area much larger than the study area. We obtained 159 independent photo sequences for bears (28 marked, 131 unmarked), 34 photo sequences of bobcats (9 marked and 25 unmarked), 48 photo sequences of cougars (24 marked, 24 unmarked), and 479 photo sequences of coyotes (55 marked, 424 unmarked; Table 1). Scat detection dogs located 86 bear scats, 95 bobcat scats, 17 cougar scats, and 772 coyote scats. Of these, we successfully genotyped 43 bear scats out of 72 attempts (60% success rate), 86 bobcat scats out of 95 attempts (91% success rate), 15 cougar scats out of 17 attempts (88% success rate), and 201 coyote scats out of 216 attempts (93% success rate). After removing duplicate scats from the same individual in the same grid cell, we had 40 bear scats from 31 individuals, 68 bobcat scats from 32 individuals, 13 cougar scats from 7 individuals, and 165 coyote scats from 83 individuals (Table 1). Maps of spatially-explicit detection data for camera and genetic analyses are provided as supplementary information (Appendix S3: Figures S1–S4).

### Telemetry Data

Across all species and methods, the inclusion of GPS data resulted in a mean decrease in the posterior modes for animal densities of 25.4% (Fig. 4, Appendix S4: Tables S1–S8). For all methods except spatial count models, including GPS data led to a mean decrease in the coefficients of variation of 18.6% (Fig. 5, Appendix S4: Tables S1–S8). But in the unmarked models, including GPS data increased the coefficients of variation by an average of 154.5% (Fig. 5, Appendix S4: Tables S1–S8). However, very low SC+GPS density estimates were responsible for the increases in the coefficients of variation in those models—standard deviations were actually smaller in SC models including GPS data than SC models lacking GPS data. Because models containing GPS data were more precise, likely because GPS data provided valuable information aiding in the identifiability of detection parameters, we consider them more reliable than models that do not include telemetry data.

**Fig. 4:**
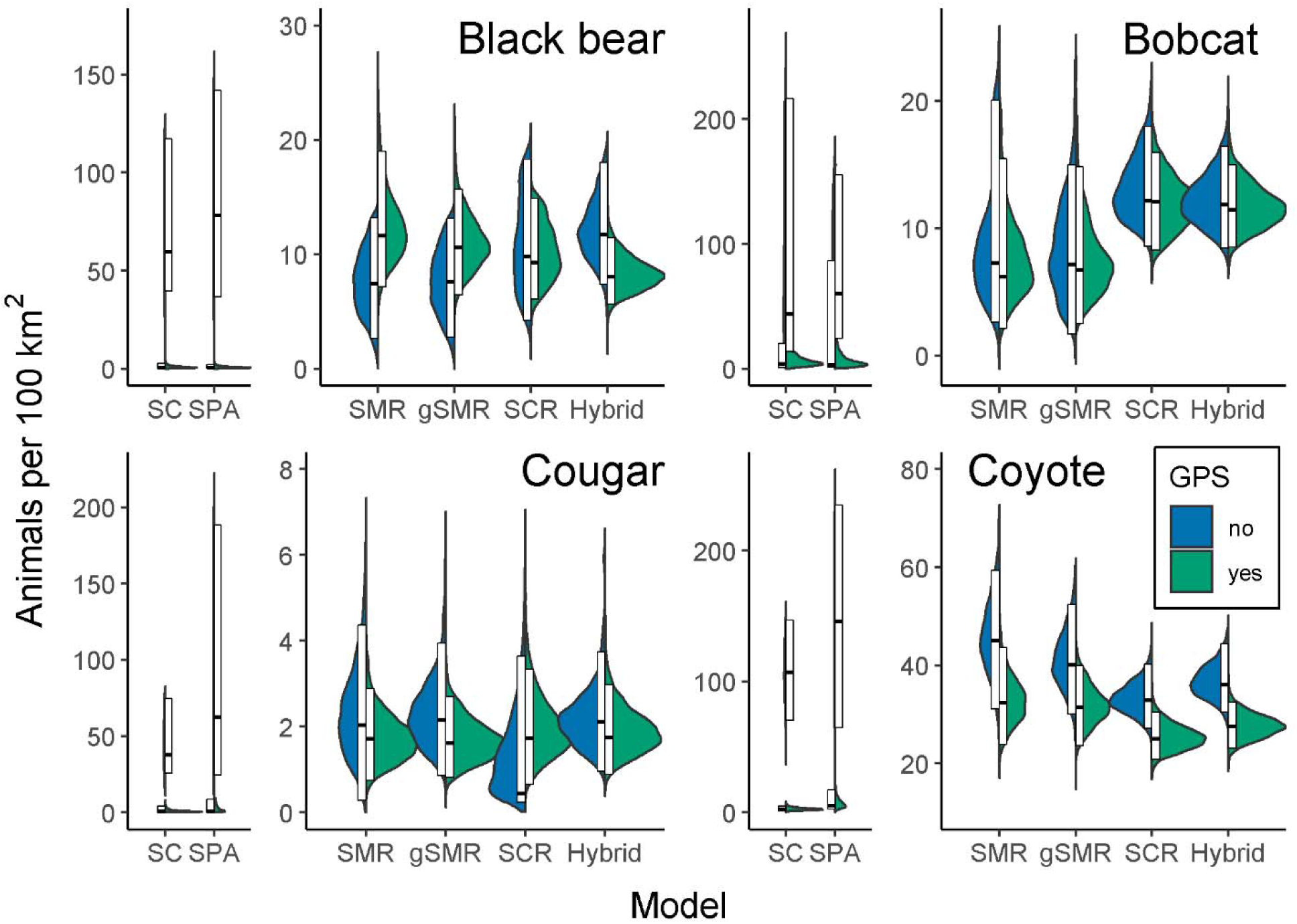
Split violin plots displaying posterior distributions for the densities of black bears, bobcats, cougars, and coyotes across a suite of methods for base models (i.e. those without sex or behavioral-specific detection parameters). Within each panel, models are ordered in terms of increasing data requirements with the most data-hungry models on the right. For each model type, the left (blue) half of the violin excludes GPS telemetry data and the right (green) half includes GPS data from a telemetered subset of individuals of each species. The white boxes display the 95% highest posterior density interval, and the middle black bar indicates the posterior mode. Densities are presented as number of animals per 100 km^2^. Note that the spatial count and spatial presence-absence models for each species required a different y-axis scale than the other model types. SC = spatial count, SPA = spatial presence absence, SMR = conventional spatial mark-resight, gSMR = generalized spatial mark-resight, SCR = spatial capture-recapture, and hybrid = combined gSMR+SCR.

**Fig. 5:**
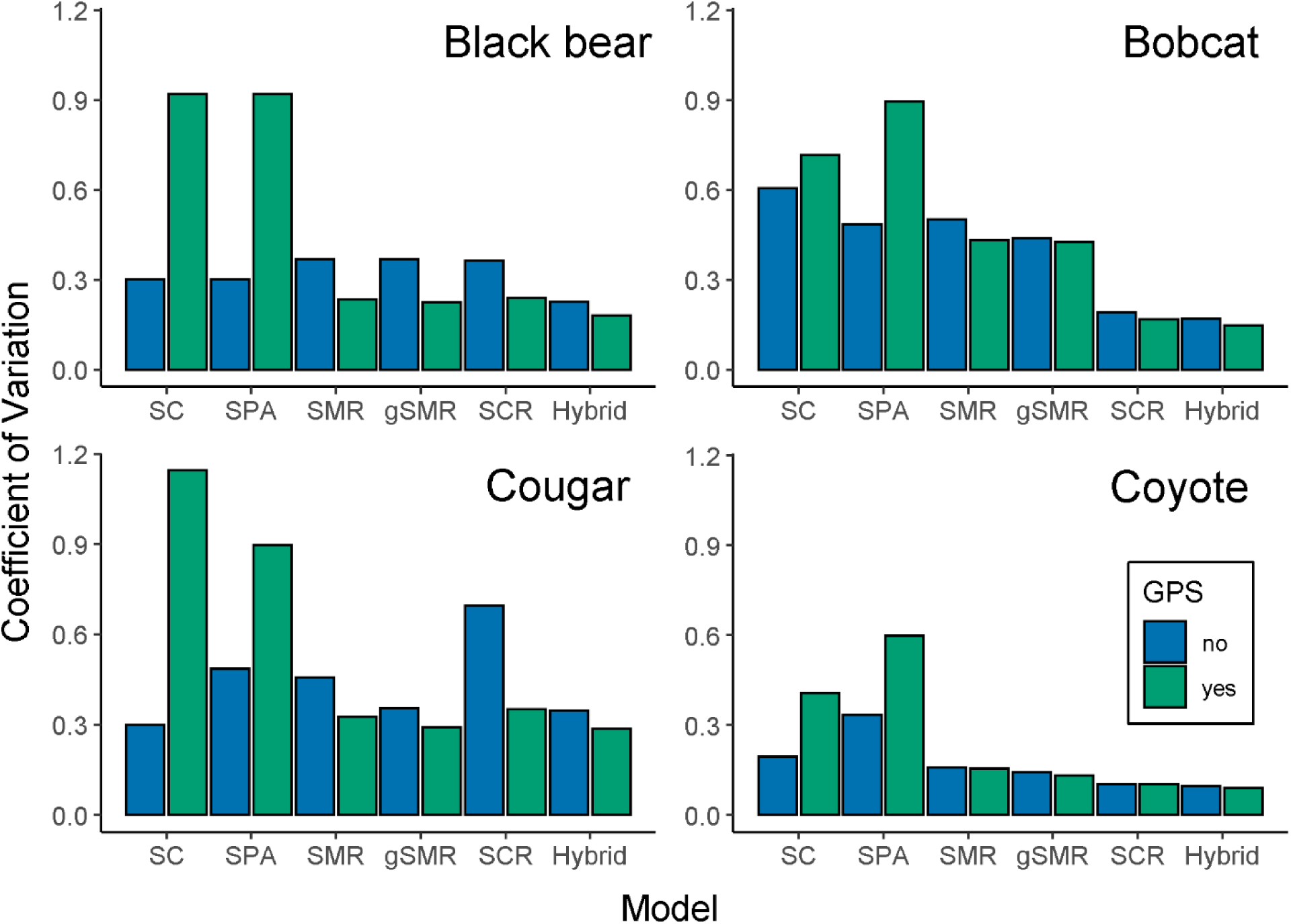
Coefficients of variation for density estimates for bears, bobcats, cougars, and coyotes (top left, clockwise). For each model, the left (blue) bar excludes GPS data and the right (green) bar includes GPS data from a telemetered subset of individuals of each species. The coefficient of variation was calculated as the posterior standard deviation divided by the posterior mean of animal density. Note that the sample size for the cougar SCR models was small (*N* = 13 scats) and likely caused the relative imprecision for those models.

### Spatial Capture-Recapture

We report SCR+GPS estimates as the benchmark to which the other methods are compared because they have the strictest requirements of individual identifiability and thus should yield the most robust estimates. The base SCR+GPS estimate of the posterior mode (and 95% BCI) for black bears was 9.9/100 km^2^ (6.1–14.9 /100 km^2^); for bobcats was 12.1/100 km^2^ (8.4–16.2/100 km^2^), for cougars was 1.7/100 km^2^ (0.9–3.6/km^2^); and for coyotes was 25.0/100 km^2^ (21.0–30.9/100 km^2^) (Fig. 4, Appendix S4: Tables S1–S8).

### Spatial Count

Spatial count models produced highly different results depending on whether telemetry data were included (Fig. 4; Appendix S4: Tables S1–S8). When GPS data were not included, densities from SC models were on average 773.9% higher than SCR+GPS estimates (Fig. 4, Fig. 6, Appendix S4: Tables S1–S8). However, when GPS data were included, densities from SC models were on average 84.4% lower than SCR+GPS estimates (Fig. 4, Fig. 6, Appendix S4: Tables S1–S8). Coefficients of variation for SC and SC+GPS models were 89.9% or 290.8% greater than base SCR+GPS, respectively (Fig. 5, Appendix S4: Tables S1–S8).

**Fig. 6:**
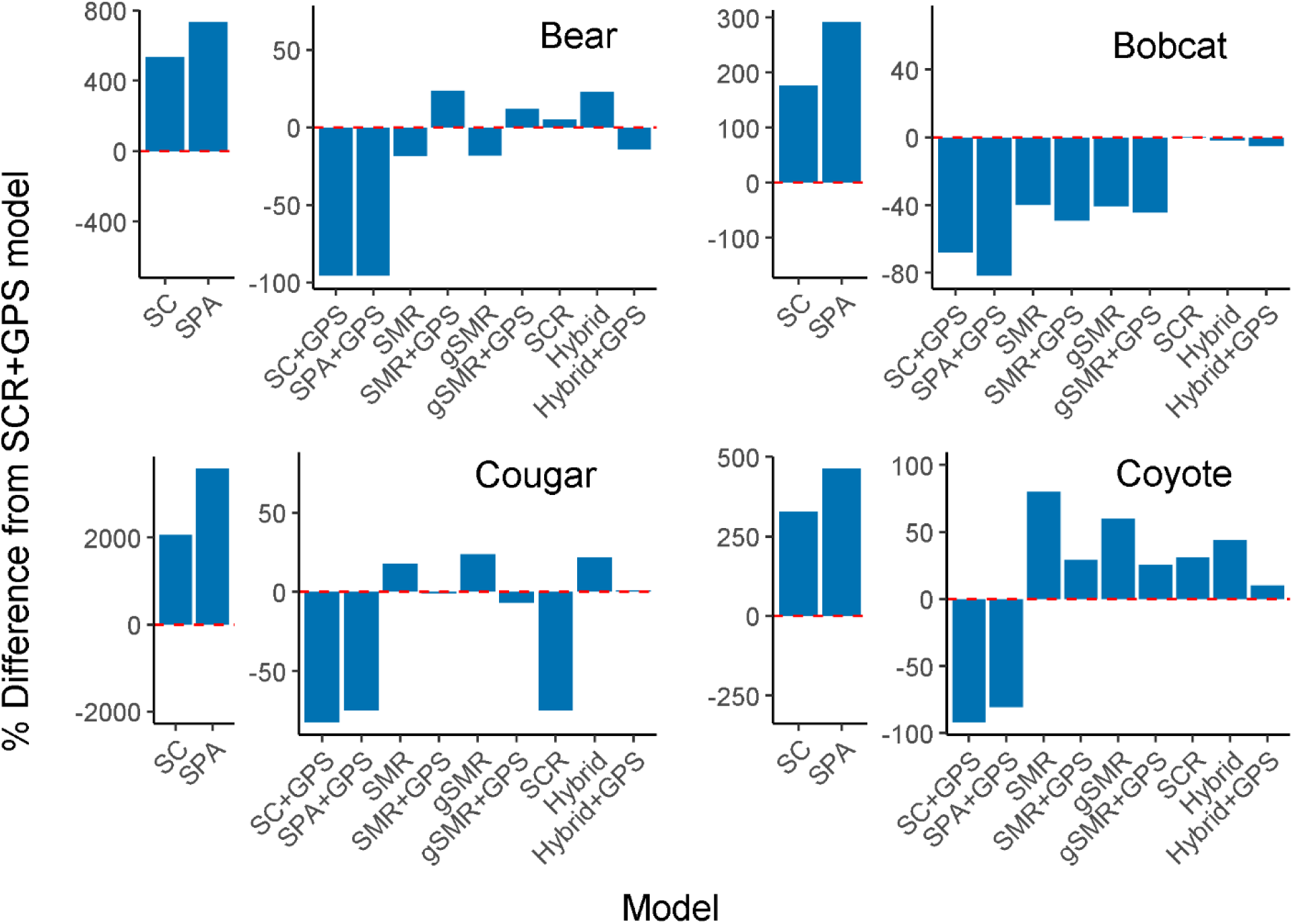
Percent difference in the posterior mode of animal density from a suite of base models compared to the base SCR+GPS (spatial capture-recapture including telemetry data) model for black bears, bobcats, cougars, and coyotes. Note that the spatial count (SC) and spatial presence-absence (SPA) models for each species required a different y-axis scale than the other model types. The dotted red line refers to no difference from the base SCR+GPS model; bars above the line indicate higher estimates and bars below the line indicate lower estimates than the base SCR+GPS model. SMR = conventional spatial mark-resight, gSMR = generalized spatial mark-resight, and Hybrid = the combined SCR+gSMR model.

### Spatial Presence Absence

Spatial presence-absence models were even more inconsistent with base SCR+GPS than were SC models. Animal densities were on average 1,267% higher or 83% lower than SCR+GPS depending on whether telemetry data were included or not (Fig. 4, Fig. 6, Appendix S4: Tables S1–S8). Coefficients of variation were 125.5% higher than the SCR+GPS model when telemetry was not included, and 470.1% higher with telemetry included (Fig. 5, Appendix S4: Tables S1– S8).

### Base Spatial Mark-Resight vs Base Spatial Capture-Recapture

Averaged over all species, base SMR+GPS models estimated animal densities that were only 0.67% lower than base SCR+GPS models (Fig. 4, Fig. 6, Appendix S4: Tables S1–S8). However, coefficients of variation were 51.8% higher in base SMR+GPS models than base SCR+GPS models (Fig. 5, Appendix S4: Tables S1–S8).

### Base Spatial Mark-Resight vs Base Generalized Spatial Mark-Resight

Across all models and species, adding the marking process to base SMR models (i.e. SMR vs gSMR) decreased estimates of animal density by an average of only 2.2% (Fig. 4, Fig. 6, Appendix S4: Tables S1–S8). However, the coefficients of variation were on average 9.8% lower in base gSMR models than base SMR models (Fig. 5, Appendix S4: Tables S1–S8). Averaged across all species, base gSMR+GPS models were 3.6% closer to the base SCR+GPS estimates than were base SMR+GPS estimates.

### Base Spatial Capture-Recapture vs Base Hybrid

The base Hybrid+GPS models estimated, on average, densities that were only 2.1% lower than base SCR+GPS models (Fig. 4, Fig. 6, Appendix S4: Tables S1–S8). But the coefficients of variation were on average 16.0% lower in the base Hybrid+GPS models than the base SCR+GPS models (Fig. 5, Appendix S4: Tables S1–S8).

### Post-hoc Analyses

The notable differences in estimates across certain base model types compelled us to conduct post-hoc analyses to attempt to explain unmodeled sources of individual heterogeneity and more strictly adhere to important model assumptions. Thus, for every species we fit an additional series of models with 1) sex-specific detection and spatial scale parameters (for all except unmarked models), 2) a reduced sampling period for all methods using camera data (70 days vs 140 days), and 3) for coyotes only, models with behavioral status-specific (i.e. resident vs transient, Appendix S5) detection and spatial scale parameters (transient coyotes in our study used home ranges that were 10–20 times the area of resident coyotes home ranges). For each of the new model types, we calculated the percent differences in posterior modes of animal densities between SCR+GPS models and SMR+GPS, gSMR+GPS, or hybrid+GPS models to assess whether the consistency of results improved. We also assessed how the coefficients of variation changed between the base and post-hoc models.

### Sex-specific Models

Including sex-specific detection and spatial scale parameters increased the SCR+GPS estimates of animal density by 4.4% averaged across species. The change in agreement between sex-specific gSMR+GPS and sex-specific SCR+GPS posterior modes of animal density relative to the same comparisons for the base models varied by species. By including sex, the difference between gSMR+GPS and SCR+GPS estimates was reduced by 5.5 percentage points (from 11.7% to 6.2%) and 10.8 percentage points (from 44.4% to 33.6%) for black bears and bobcats, respectively, and was only increased by 2.3 percentage points (from 6.9% to 9.1%) and 0.1 percentage points (25.8% to 25.9%) for cougars and coyotes (Fig. 7–8, Appendix S6: Tables S1– S8). The sex-specific SCR+GPS models had coefficients of variation that were 9.4% higher than the base SCR+GPS models when averaged across species. For gSMR+GPS, coefficients of variation were 15.3% higher in the sex-specific models than the base gSMR+GPS estimates (Fig. 9).

**Fig. 7:**
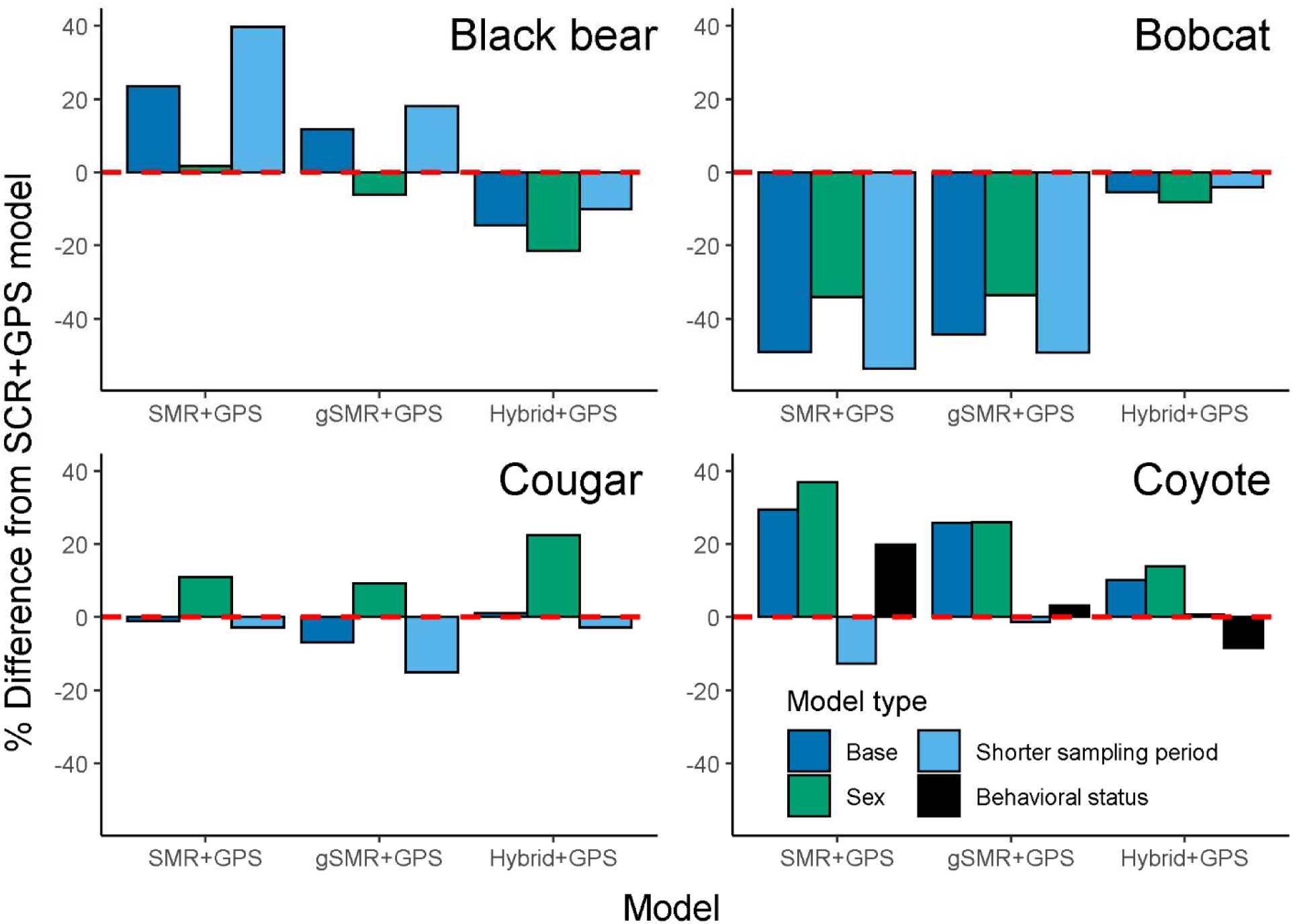
Comparison of SMR+GPS, gSMR+GPS, and Hybrid+GPS density estimates from base models and a set of post-hoc models that were fit to attempt to more fully adhere to model assumptions and/or explain more heterogeneity in detection rate in relation to SCR+GPS density estimates. Bars indicate the percent difference in the posterior mode of animal density compared to genetic SCR+GPS estimates for base models, sex-specific models (i.e. those in which detection and spatial scale parameters were allowed to vary by sex), and reduced sampling period models (reduced from 140 days to 70 days of camera sampling) for black bears, bobcats, cougars, and coyotes. For coyotes, an additional model type with detection and spatial scale parameters varying by behavioral status (i.e. transient vs resident) is compared. The horizontal, dotted red line indicates no difference from the SCR+GPS model; bars above the line indicate higher estimates and bars below the line indicate lower estimates than the SCR+GPS model. Comparisons are made with respect to sex-specific SCR+GPS estimates for ‘sex’ models, behavioral status-specific SCR+GPS estimates for ‘behavioral status’ models, and base SCR+GPS estimates otherwise.

**Fig. 8:**
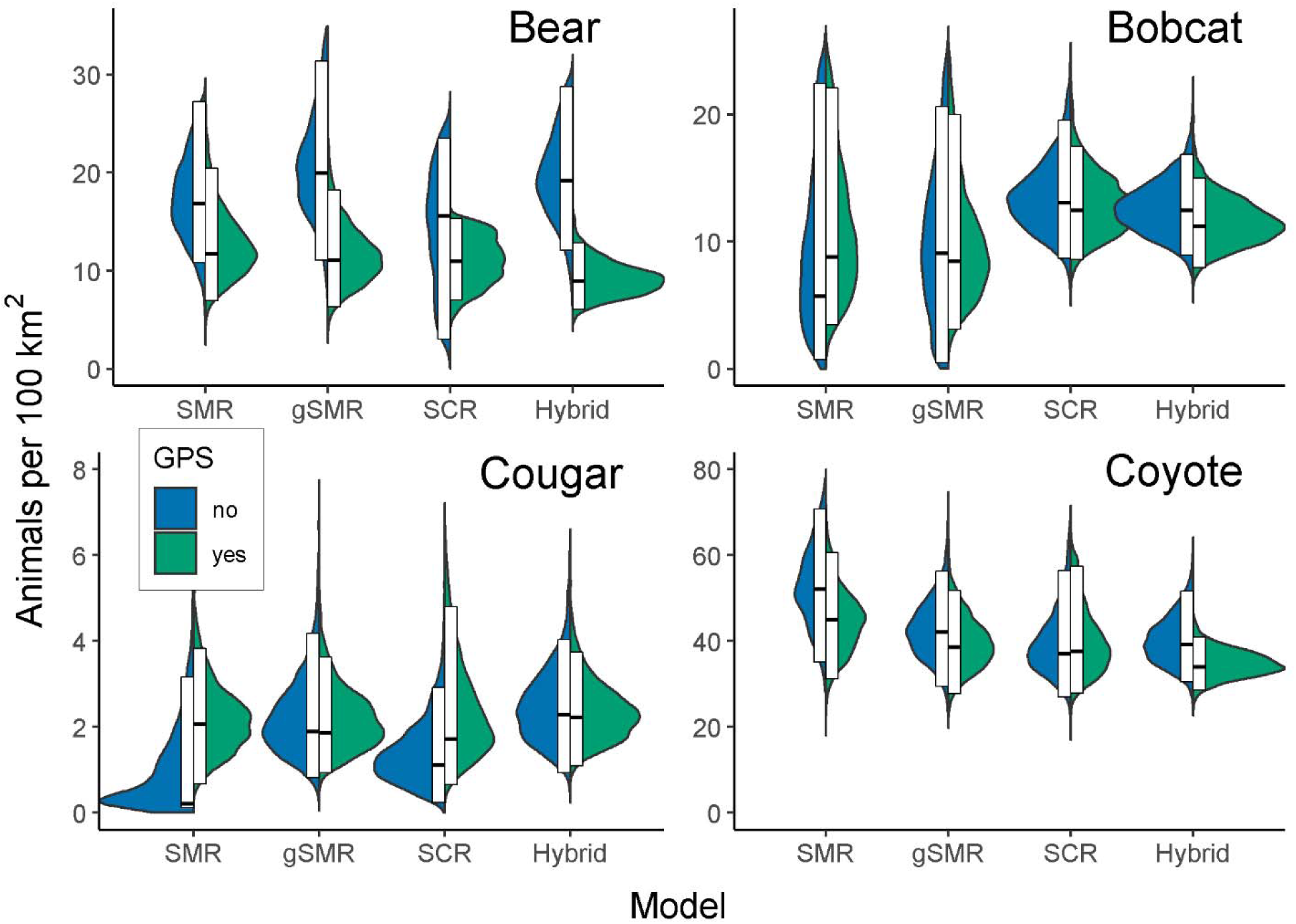
Split violin plots displaying posterior distributions for animal density estimates for sex-specific models (bobcat, bear, cougar) and behavioral specific models (resident vs transient status in coyotes). For each model type, the left (blue) half of the violin excludes GPS telemetry data and the right (green) half includes GPS data from a telemetered subset of individuals of each species. The white boxes display the 95% highest posterior density interval, and the middle black bar indicates the posterior mode. Densities are presented as number of animals per 100 km^2^. SMR = conventional spatial mark-resight, gSMR = generalized spatial mark-resight, SCR = spatial capture-recapture, and hybrid = combined gSMR+SCR.

**Fig. 9:**
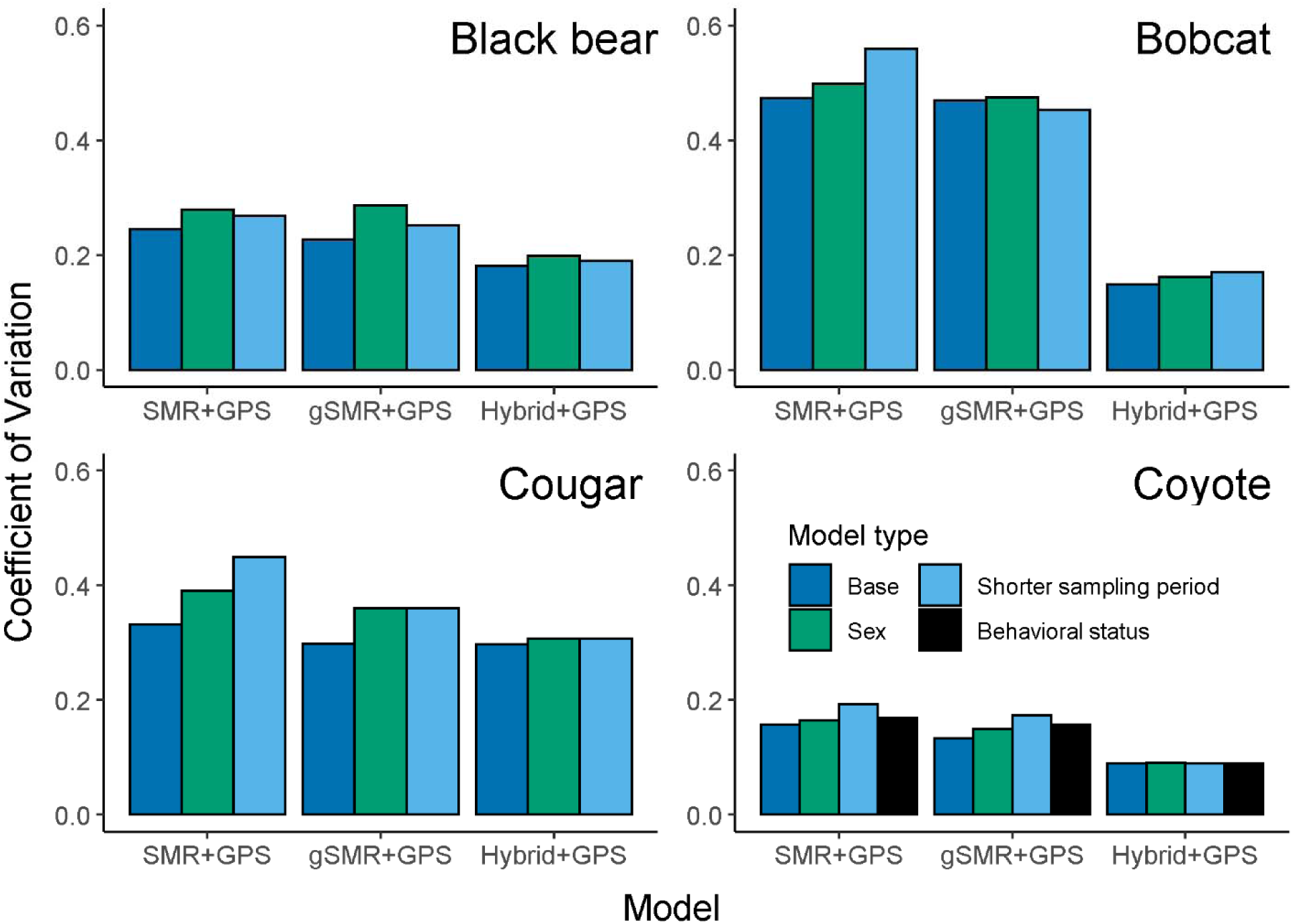
Coefficients of variation (defined as the standard deviation divided by the posterior mean) displaying the relative precision of density estimates using SMR+GPS, gSMR+GPS, and Hybrid+GPS for black bears, bobcats, cougars, and coyotes for base models, sex-specific models, and reduced sampling period models. For coyotes, an additional model type with detection and spatial scale parameters varying by behavioral status is presented.

### Reduced Sampling Period Models

When the duration of camera sampling was reduced in half for gSMR+GPS models, the estimates of animal densities were closer to the base SCR+GPS estimates for coyotes only. For that species, the percent difference in the posterior mode for animal density was reduced by 25.7 percentage points (from 25.8% to 0.1%; Fig. 7, Appendix S7: Tables S7–S8) when the reduced sampling period gSMR+GPS estimates were compared to the base SCR+GPS estimates. However, the percent difference in animal density was increased by 6.7 percentage points (from 11.7% to 18.5%), 4.9 percentage points (from 44.4% to 49.3%), and 8.1 percentage points (from 6.9% to 15.0%) for bears, bobcats, and cougars, respectively (Fig. 7, Appendix S7: Tables S1– S6). Averaged across species, the coefficients of variation for gSMR+GPS models were 13.7% higher in the reduced sampling period models than the base gSMR+GPS models (Fig. 9).

### Behavioral Status Models for Coyotes

Allowing detection and spatial scale parameters to vary by the behavioral status (i.e. resident vs transient as determined from GPS data; Appendix S5) of coyotes increased the estimates of density for models based on both camera and genetic data (Appendix S8: Tables S1–S2). However, incorporating behavioral status into models reduced the percent difference of coyote density between gSMR+GPS and SCR+GPS by 22.6 percentage points (from 25.8% to 3.16%; Fig. 7–8, Appendix S8: Tables S1–S2) relative to the same comparison for base models.

## Discussion

Many applications in ecology depend on accurate estimates of animal density. Despite tremendous advancements in wildlife monitoring technology (e.g. remote cameras), molecular methods (e.g. genotyping), and statistics (e.g. models accounting for imperfect detection), estimating population density can still be a formidable task. Using base models (i.e. those not considering variation in detection parameters induced by sex or behavioral status), we found that gSMR+GPS models based on resighting marked animals on cameras produced density estimates consistent with genetic SCR+GPS models for bears (posterior mode was 11.7% higher for gSMR+GPS than SCR+GPS) and cougars (gSMR+GPS was 6.9% lower than SCR+GPS). However, there were greater discrepancies in base models for bobcats for which the posterior mode of gSMR+GPS estimates were 44.4% lower than the SCR+GPS results and for coyotes for which gSMR+GPS estimates were 25.8% higher than the SCR+GPS results. For both species, the posterior mode of the SCR models was contained in both the 95% HPDI and BCI of SMR models, but the converse was not true (Fig. 4, Appendix S4: Tables S3 and S7).

Although base gSMR models and base SCR models produced qualitatively similar density estimates, the moderate differences in results for bobcats and coyotes may be due to challenges of simultaneously camera trapping for multiple species with distinct habitat preferences, home range sizes, and life history strategies. We attempted to refine estimates by fitting additional post-hoc models to explain more variation in heterogeneity of detection rate using individual covariates on detection parameters for sex, and, for coyotes only, behavioral status to consider resident and transient states. After controlling for these important sources of variation, we found much greater concordance in estimates between camera-based gSMR and genetic SCR (Fig. 8).

We also fit models using half the sampling occasions for camera analyses to assess whether a lack of demographic closure could have driven the discrepancies between analyses based on camera versus genetic data. While the available data indicated that demographic closure was not grossly violated in this study (one marked cougar died during the time of camera sampling), it is nonetheless possible that other deaths or dispersal occurred which would cause our estimates to be biased high. If, however, recruitment occurred and juveniles were not detected then our estimates could be biased low. But because the species considered here are generally long-lived and our sampling was designed to miss the birth pulse, we believe the population was reasonably closed during our study.

Base SCR+GPS estimates were 44% higher than base gSMR+GPS estimates for bobcats, but this was reduced to 33% by including sex-specific detection parameters. This modest improvement in consistency was likely driven by sex-specific differences in home range size, which induced heterogeneity in detection rate that was previously unaccounted for. However, a moderate discrepancy in estimates remained. Based on GPS data, bobcats in this system used dense forest cover and our camera placements were biased away from this habitat type because they had extremely limited viewsheds that would limit photos of other species. In contrast, detection dogs readily located bobcat scats and the higher detection probability of scats may have driven the difference in results between cameras and genetics. Alternatively, the assumption of demographic closure may have been violated since bobcats can give birth nearly year-round (Winegarner and Winegarner 1982). It is possible that juvenile bobcats were detected via genetic sampling but that they were missed on cameras. Genetic techniques are currently unable to accurately determine the age of the individual from its scat, so unless scats of juveniles can be censored due to size or other morphological characteristics, recruitment has the potential to be problematic for all genetic mark-recapture studies.

For coyotes, the inconsistency between base gSMR and base SCR models likely involved unmodeled sources of individual heterogeneity in detection rates across methods as was observed by Murphy et al. (2018a) who estimated lower densities when models were constructed solely from scat data than from both scat and hair. The authors speculated that the behavioral status of coyotes (i.e. resident vs transient) led to cohorts that were disproportionately detected using one detection method over another. Specifically, they reasoned that territorial resident coyotes could be more likely to be detected from scats because of smaller home ranges and increased scent-marking with scat for territorial defense than wide-ranging transients. Our study may exhibit an analogous issue if cameras detected both resident and transient coyotes at similar rates but the detected scats disproportionately belonged to residents. Similarly, Morin et al. (2016) suggested that SCR density estimation in coyotes could be improved by modeling heterogeneity in σ and baseline detection rates arising as a result of behavioral status. Without modelling such heterogeneity, the bimodality in home range size can lead to estimates of detection and scale parameters that are averaged between the two behavioral states (Murphy et al. 2018). Transient coyotes in our study used home ranges that were 10–20 times greater than those of resident coyotes.

To explore whether failing to acknowledge these sources of heterogeneity were responsible for the inconsistencies between base gSMR and base SCR models, we conducted a post-hoc analysis where σ and baseline detection parameters were allowed to vary as a function of transient or resident behavioral status (see Appendix S5 for details). Including behavioral status increased the estimates of density from base models in both gSMR+GPS and SCR+GPS models but reduced the difference between them substantially from 26% to 3% (Fig. 7, Appendix S8: Tables S1–S2). We thus speculate that the unmodeled heterogeneity arising from the distinct behavioral states explained much of the disagreement between model types. We recommend others control for behavioral states in coyotes if GPS data are available to classify individuals into residents or transients.

Spatial count and spatial presence-absence models reliant on photos of unmarked animals produced density estimates that were very inconsistent with methods in which individuals were identifiable. For all four species, we observed the same pattern in which estimates from SC and SPA models without GPS data were higher than SCR estimates and the same models with GPS data were lower than SCR estimates. Informing σ with telemetry data did not lead to accurate enumeration of activity centers and instead falsely attributed unmarked photo detections to too few individuals. Overdispersion in the count data was probably not responsible for imprecise results in the SC models because after collapsing counts to independent binary detections and using the SPA model, the estimates became even more different from SCR results.

SC and SPA models require sufficiently close trap spacing so that spatial autocorrelation can be detected and used to identify activity centers (Chandler and Royle 2013, Ramsey et al. 2015). Poor density estimation from SC models can result from insufficient spatial structure of animal detections that prevent individual activity centers from being clearly demarcated. However, our 1 km camera grid spacing allowed for many cameras per animal home range, suggesting that these models are not robust for carnivores under sampling scenarios that are feasible to accomplish in most studies. Further, SC and SPA results can be unreliable, unstable, or suffer from convergence failure when data are sparse (Chandler and Royle 2013, Ramsey et al. 2015, Kane et al. 2015, Burgar et al. 2018a, Burgar et al. 2018b, Murphy et al. 2018b), suggesting that this will be a consistent challenge in carnivore studies using unbaited cameras within a time frame over which demographic closure can be reasonably assumed. Populations with large values of σ and/or high densities can produce biased results in SC models (Chandler and Royle 2013, Augustine et al. 2019) limiting their potential for use in many species. Including habitat covariates that influence spatial density in SC models may lead to results that are consistent with SCR (e.g. Evans and Rittenhouse 2018), but our results suggest that fully unmarked SCR models (i.e. SC and SPA) may unsuitable for density estimation of the carnivores we studied, particularly when detectors are widely spaced, count data are sparse, and auxiliary data are not available.

Previous studies have successfully estimated population densities of multiple carnivore species using camera trapping (Jiménez et al. 2017, Burgar et al. 2018a, Rich et al. 2019). Our study also suggests a multi-species monitoring approach is possible yet highlights the challenges of simultaneous density estimation of multiple species. For example, the substantial variation in home range size among carnivores requires fine-scale sampling for smaller ranges (e.g. coyotes and bobcats) while allowing for the larger extents needed for greater home range sizes (e.g. bears and cougars). This can be accomplished with either a finely-spaced grid (this study) or through cameras that are clustered with variable spacing intervals (Rich et al. 2019, Murphy et al. 2019). Similarly, assumptions of demographic closure may be challenging to meet for species that roam widely unless the extent of the study area is large enough to contain many home ranges. For example, our small survey area relative to the species with the largest home range and lowest density (e.g. only 7 distinct cougars were sampled from genetics) may have made our estimates sensitive to the assumption of demographic closure given that the death, dispersal, or recruitment of even one individual could have a substantial influence on estimates.

For all species, the inclusion of telemetry data to inform density estimation generally produced consistent results with increased precision and smaller coefficients of variation for all methods except unmarked models (Fig. 5, Appendix S4: Tables S1–S8). The gains in precision were greatest for the models/species with the sparsest data (e.g. cougar SCR model). This highlights the challenge of estimating σ when telemetry data are excluded and spatial recaptures are sparse. For example, transient coyotes with large home ranges may rarely or never be recaptured, which would falsely suggest low detectability rather than the large value for σ that is evident from telemetry data. The absence of telemetry data can thus lead to both imprecise and biased estimates of σ with consequent effects on other model parameters (i.e. λ0 or *p*0, *s_i_*, and ultimately, *N*) (Whittington et al. 2018). Improving the estimate of σ by incorporating telemetry data should aid in identifiability of other parameters and lead to greater accuracy in the models incorporating GPS data. SC and SPA models were particularly problematic because they produced biologically unrealistic density estimates that changed from very high to very low with the inclusion of telemetry data. Although telemetry data decreased precision in unmarked models, the large reduction in the density estimate when including telemetry data led to an increase in the coefficient of variation because the denominator became smaller. An inaccurate result with a low CV invites the possibility of overconfidence in an incorrect result when using unmarked models.

Models that included more information on individual identity exhibited more precise density estimates (Fig. 5), except when the number of genotyped scats was very low. In particular, the small sample of cougar scats led to SCR models that were substantially less precise than SMR and gSMR models (Fig. 5, Appendix S4: Table S5). For all species, the hybrid model containing camera, physical capture (gSMR), and genetic (SCR) data had the lowest coefficients of variation (Fig. 5) highlighting the benefit of leveraging multiple datasets even if some are sparse.

A recent movement in ecological modeling of demographic rates has sought to strengthen inference by utilizing multiple independent data sources (e.g. Besbeas et al. 2002, Schaub et al. 2007). SMR and SCR models require a substantial investment of time and money yet may still only yield sparse data. Integrating multiple sparse data sources is thus an efficient use of resources in addition to the benefits of improving estimation. Others have integrated camera and genetic data for spatially-explicit density estimation (Sollmann et al. 2013b, Clare et al. 2017, Gopalaswamy et al. 2012), but to our knowledge this has only occurred when animals on camera could always be identified to individual, so the result was a SCR model composed of two different data sources. And recently, both identified and unidentified genetic samples were combined in a SCR model using multiple observation processes (Tourani et al. 2020). These studies, however, contrast with our approach of combining SMR for a partially-marked population (i.e. those in which not all individuals can be uniquely identified) with genetic SCR data in which every sample was identifiable to individual, and sharing information via deterministic linkages between individuals that were genotyped, GPS collared, and resighted on cameras.

Ancillary data associated with genetic or mark-resight data may motivate the decision to collect one data type over another. For example, telemetry data obtained in conjunction with a mark-resight study may be useful not only to inform σ, *p*0/λ0, and the activity centers of the telemetered individuals, but also to classify an animal into a behavioral state such as resident vs transient or breeder vs nonbreeder, and these distinctions may explain important sources of individual heterogeneity in detection rate. The photographs in a mark-resight analysis may also permit identification of juveniles which can easily be censored if the researcher desires to estimate the density of the adult population segment only. While genetic sampling is unlikely to yield these sources of information, it offers other benefits such as the ability to study diet (e.g. Kartzinel et al. 2015), hormone levels (e.g. Gobush et al. 2014), and population structure or genetic diversity (e.g. Goosens et al. 2005), which if useful for other objectives may steer researchers toward that method of data collection. Genetic samples also readily yield the sex of the individual which may not be possible from camera trapping for species that are not sexually dimorphic. The cost of each data collection method may also play a factor in the decision. While both genetics and mark-resight studies may be expensive, when a segment of the population is already being monitored via telemetry devices, the extra cost to resight marked individuals using cameras may be minimal compared to the cost of a separate genetics SCR study (Whittington et al. 2018). However, in the absence of an existing effort to obtain telemetry data, genetic SCR methods are substantially more cost effective given the expense associated with capturing, collaring, and resighting animals.

Another extension to our efforts might implement spatial partial-identity models to extract even more information from the existing datasets (Augustine et al. 2018). This would allow partial genotypes (Augustine et al. 2019) or photos in which the marked status or individual identity (Augustine et al. 2018, Jiménez et al. 2019, Tourani et al. 2020) cannot be ascertained to be included in SCR estimation. However, this model extension would minimally influence our results because of the high genotyping success from scats using genotyping by amplicon sequencing (Eriksson et al. 2020) and our high success in identifying marked status or individual identity in our camera trap photos. Further, if the discrepancies between model types were driven by violations of model assumptions (e.g. demographic closure), partial identity models would likely not overcome the inherent bias and may instead only increase precision.

Knowing how much and what type of data to collect is a challenge confronted by most researchers aiming to efficiently estimate population densities. Our results from a suite of spatially-explicit models for bears, bobcats, cougars and coyotes in a natural system highlight the relative performances of models within the spatial capture-recapture family constructed from data of varying sources and resolutions. We acknowledge that in some instances, such as setting management objectives for common species, absolute population densities may not be essential and simply being able to accurately characterize relative changes in population trends may be adequate. In other cases, such as the calculation of extinction risk for imperiled species, obtaining the most accurate estimates of population size is crucial. We found that the most common method for estimating population size of terrestrial carnivores in the past five years was (spatial) capture-recapture and that relatively few studies used methods that do not require individual recognition (Appendix S1: Fig. 1). This suggests that in most cases, researchers are willing to spend the resources necessary to achieve robust results. Regardless, the method chosen must be commensurate with the objective of the study.

We conclude by suggesting that future research employing SCR or its variants to estimate the density of carnivores (1) ensures the assumptions and study design requirements of unmarked methods can strictly be met before using them, (2) controls for important sources of variation in detection probability such as those arising from sex or behavioral status, (3) incorporates GPS data when possible, and (4) considers combining multiple data sources, e.g. using the hybrid gSMR+SCR model presented here. We note that SCR models with few scats produced density estimates consistent with, but less precise than, a years-long effort to capture carnivores and resight them on camera traps. Given the inadequacy of the unmarked models and expense associated with capturing carnivores to mark them for resighting, we recommend that new studies to estimate population density of carnivores (that are not distinguishable with natural markings) first consider using genetic SCR. The efficacy of this approach will continue to increase as noninvasive genetic sampling transitions toward low-cost and high-success genotyping of SNPs by high-throughput sequencing (Eriksson et al. 2020). Finally, newer models to estimate density using unmarked animals, such as the time-to-event and space-to-event family of models (Moeller et al. 2018), provide hope for a cost-effective method to estimate population densities for multiple species. These models, however, have their own set of assumptions such as random camera placement with respect to animal movement, accurate measurement of the camera viewshed area and perfect detection of animals within that area, and in the case of the time-to-event model, accurate and independent estimates of animal movement speed. Whether those assumptions are tenable for typical field applications will require a similar test to that presented here prior to their integration into large-scale carnivore monitoring programs.

## Supporting information

density methods lit review

genotyping methods

spiderplots

base density tables

coyote behavioral status methods

sex model updates

reduced sampling model tables

behavioral status model tables

## Acknowledgments

We thank M. Bianco, K. Bowman, C. Brown, L. Carr, L. Chodelski, T. Craddock, E. Crain, C. Eckrich, M. Esquibel, S. Gillman, M. Goldman, A. Hilger, R. Jensen, C. Kidd, M. Olsen, J. Smith, A. Yancey, and J. Yancey for field support. We thank K. Cronin, S. Gillman, M. Goldman, J. Hart, V. Heilman, R. Jensen, J. Neeley, N. Owen, S. Muñoz for photo tagging support. We thank J. Allen for lab work. We thank R. Chandler and B. Augustine for coding assistance. We thank R. Chandler and an anonymous reviewer for helpful comments on an earlier draft. We thank B. Dick, R. Kennedy, B. Naylor, and D. Rea and the USDA Forest Service Pacific Northwest Research Station for logistical support. Funding and/or field equipment was provided from Oregon Hunter’s Association, Oregon Trapper’s Association, Wildlife Restoration Act, Oregon Department of Fish and Wildlife, USDA Forest Service, and Oregon State University.

## Notes

https://github.com/taaltree/SCR

### Competing Interest Statement

The authors have declared no competing interest.

### Summary of Updates

We now additionally use specific models (bobcat, bear, cougar) and behavioral specific models (resident vs transient status in coyotes) to determine post hoc whether model variants converge to a more similar answer when accounting for additional unmodeled heterogeneity. There are additional stylistic changes throughout.

