## Supplementary material for "Evaluating and integrating spatial capture-recapture models with data of variable individual identifiability": density methods lit review

**Appendix S1**: Literature review of density estimation methods for carnivores, 2015–2020.

We queried Google Scholar between 18 and 20 March 2020 to attain a sample of the methods used to estimate carnivore population densities using search terms “carnivore density estimation” and “carnivore abundance estimation.” For each search term we reviewed the first 250 records only. We only utilized studies that estimated the absolute population density or abundance (i.e. not relative measures or indices of abundance or density) of terrestrial carnivores between 2015 and 2020. The methods in each study were classified into one of nine categories: spatial count (SC; Chandler and Royle 2013), spatial mark-resight (SMR; Sollmann et al. 2013), spatial capture-recapture (SCR; Royle et al. 2013), capture-recapture (e.g. Otis et al. 1978, Nichols 1992), mark-resight (e.g. White 1996, McClintock and White 2009), Royle-Nichols model (Royle and Nichols 2003), random encounter model (Rowcliffe et al. 2008), N-mixture model (Royle 2004), distance sampling (Buckland et al. 1993), and other.

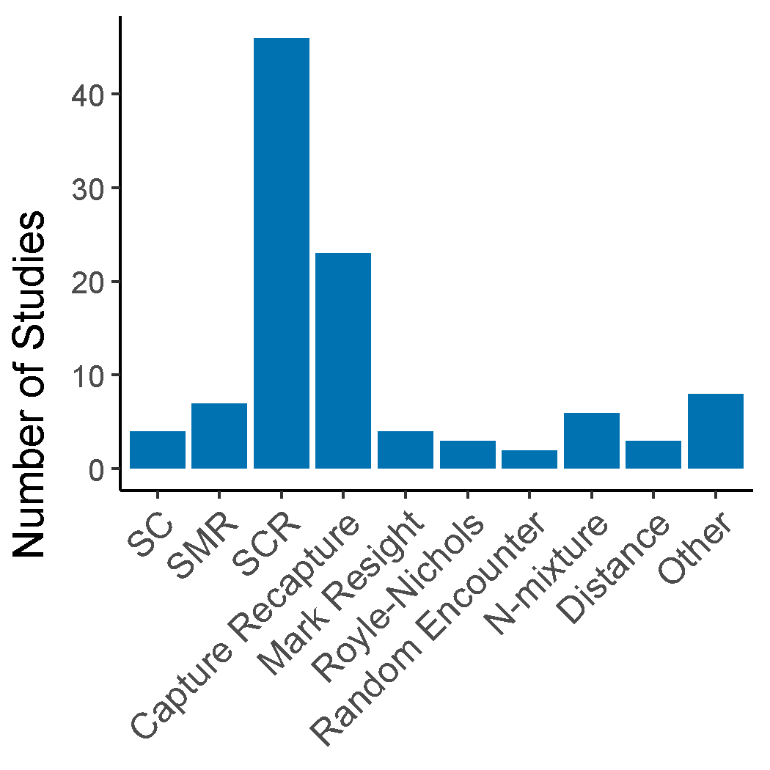

**Figure S1**: The frequency of methods used for the density estimation of terrestrial carnivores between 2015 and 2020 (N = 88 studies reporting methods 106 times).
