## Supplementary material for "Evaluating and integrating spatial capture-recapture models with data of variable individual identifiability": genotyping methods

**Appendix S2**: Single Nucleotide Polymorphism (SNP) Discovery and Selection.

We designed locus-specific primers based on previously published SNP positions and flanking regions for coyotes (Monzón et al. 2014, Monzón 2014) and cougars (Fitak et al. 2015). For black bears we used SNPs identified from RAD-seq data in Puckett et al. (2015). We filtered for samples collected in Oregon (N = 5) and calculated allele frequencies using Plink (Purcell et al. 2007). We mapped the positions of the SNPs to the polar bear genome (Liu et al. 2014) and extracted 100 bp flanking region on either side of the SNPs using Geneious (https://www.geneious.com).

Since no SNPs were published for bobcats we aligned SNPs from European wildcat (Nussberger and Wandeler 2014) and Iberian lynx (Abascal et al. 2016) to Sequence Read Archive (SRA) data from bobcats in California (Fraser et al. 2018). We used BLAT to align the sequences containing the SNPs (101 bp) from both species to one individual bobcat. The sequences that matched were then mapped to each individual bobcat (N = 52) with BWA-MEM (Li and Durbin, 2009), followed by conversion and sorting using SAMtools (Li et al. 2009). We used the RealignerTargetCreator and IndelRealigner in the Genome Analysis Toolkit (GATK) (McKenna et al. 2010) to realign deletions or insertions. For variant calling we first generated a vcf using SAMtools mpileup, filtered out low quality SNPs using ‘QUAL>10’, and SNPs with missing information in many individuals (‘--max-missing 0.85’.). We selected SNPs with 2 alleles, Phred quality score > 20, non-significant deviation from Hardy-Weinberg Equilibrium, and minimum allele frequency > 0.1 (See Table S1 for total number of SNPs genotyped). SNP primer design, optimization and genotyping protocol followed methods outlined in Eriksson et al. (2020).

**Table S1**: Total number of Single Nucleotide Polymorphisms (SNPs) genotyped, (probability of identity) PID and probability of identity of siblings (PID-sibs) values for each species.

| Species | Number of SNPs | PID | PID(sibs) |
| --- | --- | --- | --- |
| Bobcat | 24 | 0.0000002 | 0.0004 |
| Black bear | 26 | 0.0000000007 | 0.00001 |
| Cougar | 24 | 0.0000008 | 0.0007 |
| Coyote | 26 | 0.000001 | 0.0009 |

genotyping using high-throughput amplicon sequencing. Molecular Ecology Resources

https://doi.org/10.1111/1755-0998.13208.

Fitak, R. R., A. Naidu, R. W. Thompson, and M. Culver. 2015. A new panel of SNP markers for the individual identification of North American Pumas. Journal of Fish and Wildlife Management 7:13–27.

Fraser, D., A. Mouton, L. E. K. Serieys, S. Cole, S. Carver, S. Vandewoude, et al. 2018. Genome-wide expression reveals multiple systemic effects associated with detection of anticoagulant poisons in bobcats (*Lynx rufus*). Molecular Ecology 27:1170–1187.

Li, H., and R. Durbin. 2009. Fast and accurate short read alignment with Burrows – Wheeler transform. Bioinformatics 25:1754–1760.

Li, H., B. Handsaker, A. Wysoker, T. Fennell, J. Ruan, N. Homer, et al. 2009. The Sequence Alignment/Map format and SAMtools. Bioinformatics 25:2078–2079.

Liu, S., E. Lorenzen, M. Fumagalli, B. Li, K. Harris, Z. Xiong, et al. 2014. Population genomics reveal recent speciation and rapid evolutionary adaptation in polar bears. Cell 157:785–794.

McKenna, A., M. Hanna, E. Banks, A. Sivachenko, K. Cibulskis, A. Kernytsky, et al. 2010. The Genome Analysis Toolkit: A MapReduce framework for analyzing next-generation DNA sequencing data. Genome Research 20:1297–1303.

Monzón, J., R. Kays, and D. E. Dykhuizen. 2014. Assessment of coyote-wolf-dog admixture using ancestry-informative diagnostic SNPs. Molecular Ecology 23:182–197.

Monzón, J. 2014. First regional evaluation of nuclear genetic diversity and population structure in northeastern coyotes (*Canis latrans*). F1000Research 3:1–15.

Nussberger, B., and P. Wandeler. 2014. A SNP chip to detect introgression in wildcats allows accurate genotyping of single hairs. European Journal of Wildlife Research 60:405–410.

Puckett, E. E., P. D. Etter, E. A. Johnson, and L. S. Eggert. 2015. Phylogeographic analyses of American black bears (*Ursus americanus*) suggest four glacial refugia and complex patterns of postglacial admixture. Molecular Biology and Evolution 32:2338–2350.

Purcell, S., B. Neale, K. Todd-Brown, L. Thomas, M. A. R. Ferreira, D. Bender, et al. 2007. PLINK: A tool set for whole-genome association and population-based linkage analyses. The American Journal of Human Genetics 81:559–575.
