## Supplementary material for "Evaluating and integrating spatial capture-recapture models with data of variable individual identifiability": spiderplots

**Appendix S3**: Maps of detection data for spatial density models of black bears, bobcats, cougars, and coyotes.

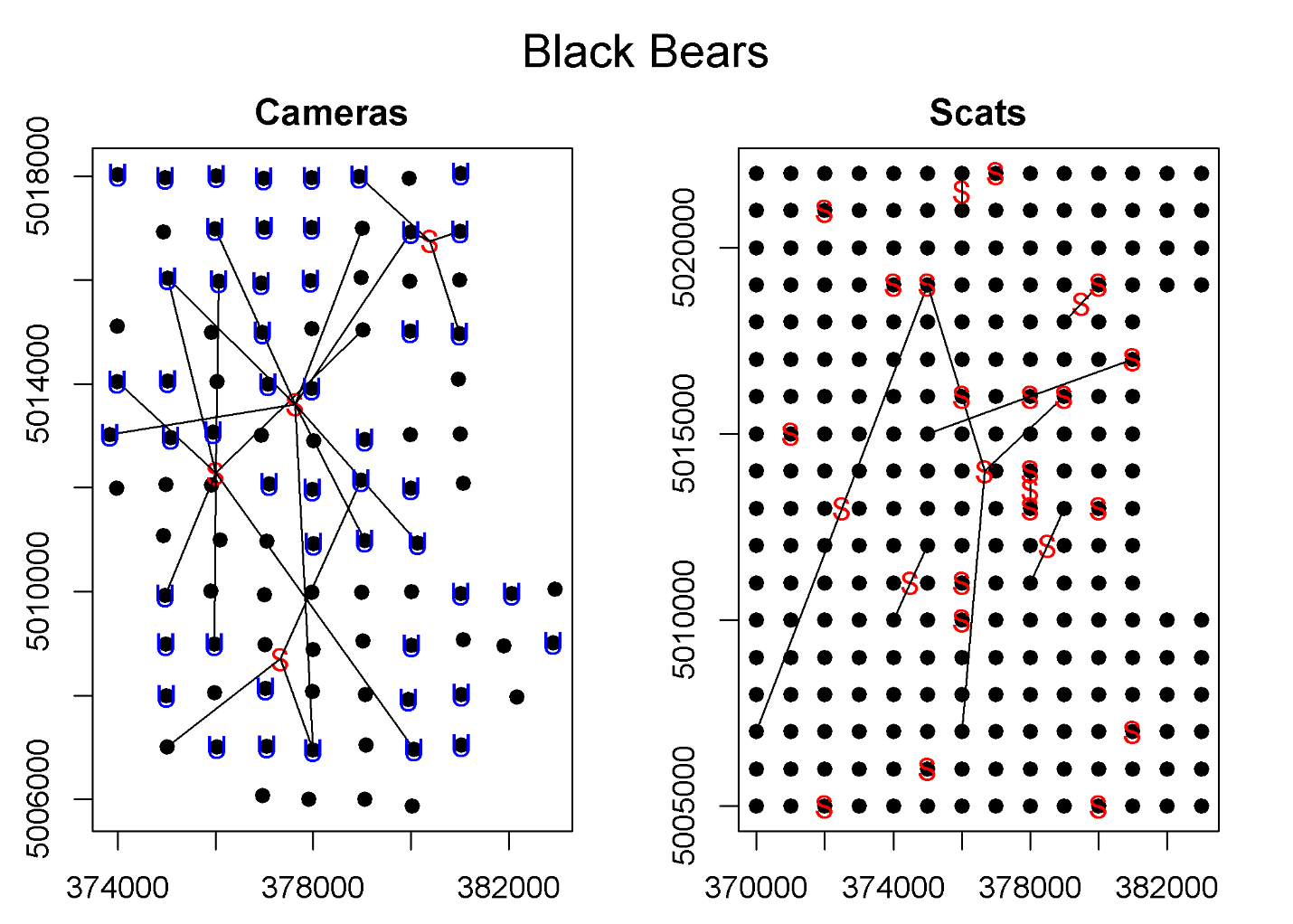

**Figure S1**: “Spiderplots” showing locations of black bear detections from remote cameras (left panel) or scats located by detector dogs and genotyped (right panel). Black dots represent the camera location (left panel) or the center of each grid cell (right panel). In both panels, the detections for a given individual are connected by black lines and the red “S” displays the centroid of the detections for that individual. In the left panel, the blue “U” designates cameras that detected unmarked black bears.

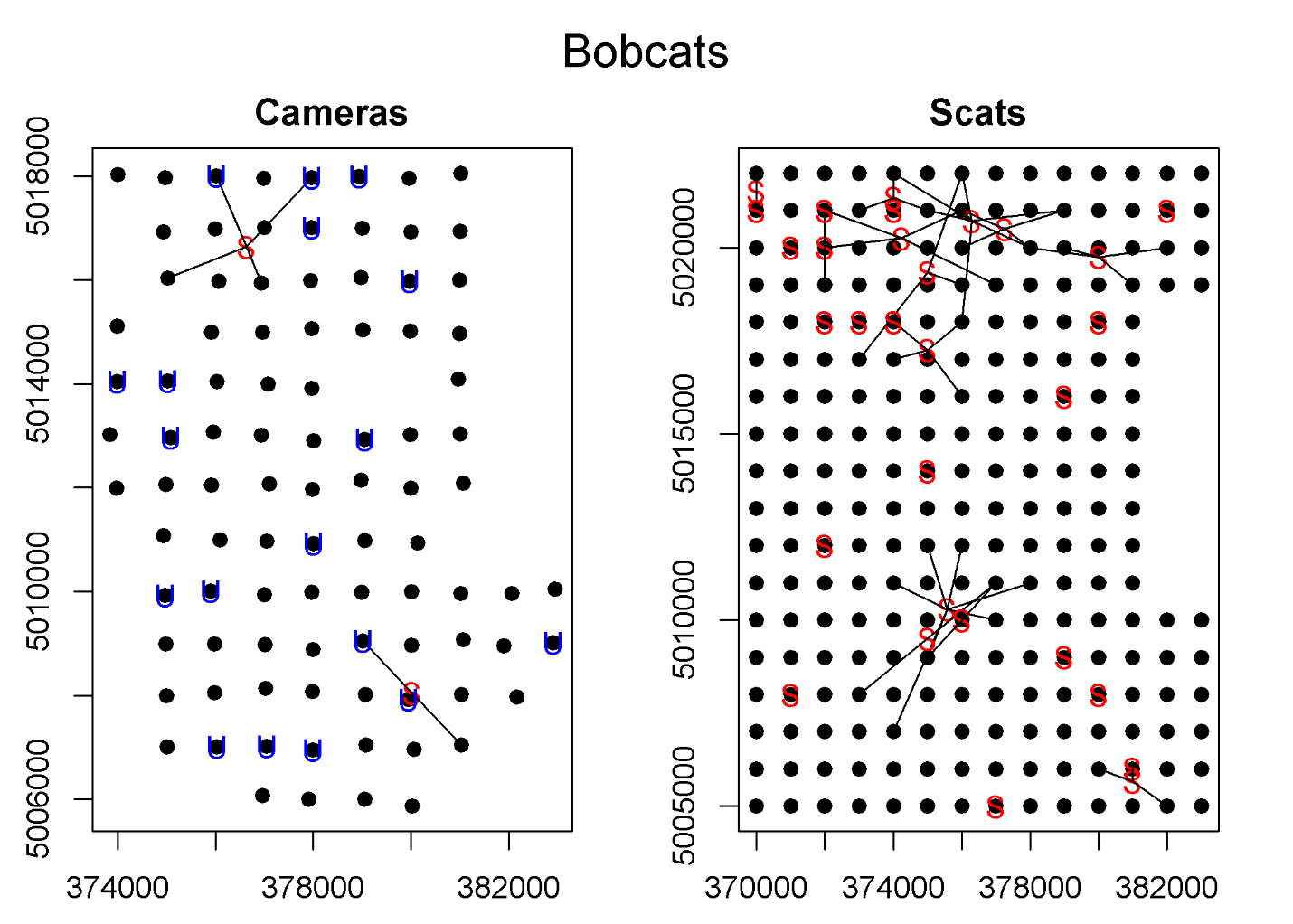

**Figure S2**: “Spiderplots” showing locations of bobcat detections from remote cameras (left panel) or scats located by detector dogs and genotyped (right panel). Black dots represent the camera location (left panel) or the center of each grid cell (right panel). In both panels, the detections for a given individual are connected by black lines and the red “S” displays the centroid of the detections for that individual. In the left panel, the blue “U” designates cameras that detected unmarked bobcats.

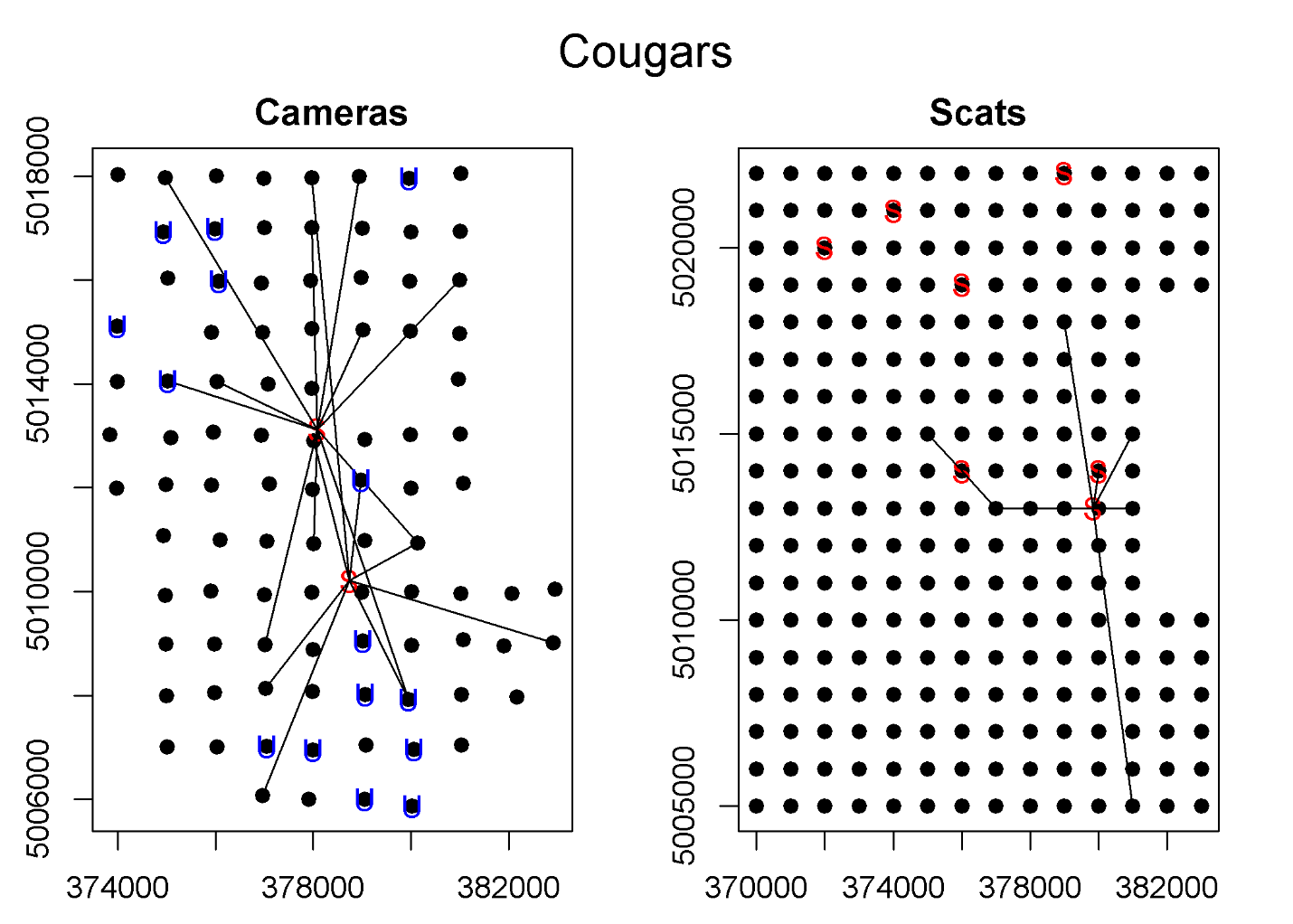

**Figure S3**: “Spiderplots” showing locations of cougar detections from remote cameras (left panel) or scats located by detector dogs and genotyped (right panel). Black dots represent the camera location (left panel) or the center of each grid cell (right panel). In both panels, the detections for a given individual are connected by black lines and the red “S” displays the centroid of the detections for that individual. In the left panel, the blue “U” designates cameras that detected unmarked cougars.

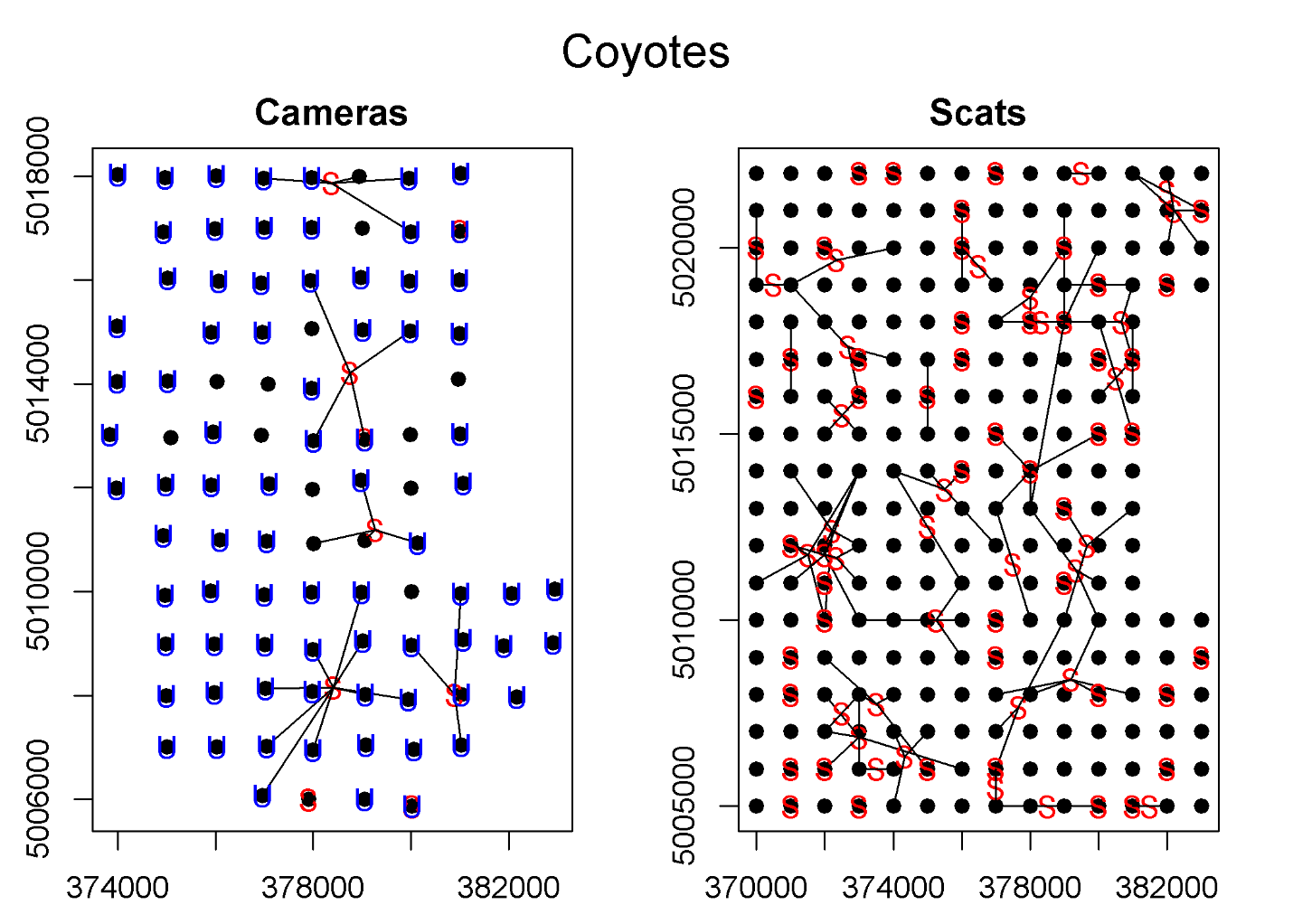

**Figure S4**: “Spiderplots” showing locations of coyote detections from remote cameras (left panel) or scats located by detector dogs and genotyped (right panel). Black dots represent the camera location (left panel) or the center of each grid cell (right panel). In both panels, the detections for a given individual are connected by black lines and the red “S” displays the centroid of the detections for that individual. In the left panel, the blue “U” designates cameras that detected unmarked coyotes.
