## Supplementary material for "Evaluating and integrating spatial capture-recapture models with data of variable individual identifiability": coyote behavioral status methods

**Appendix S5**: Description of conventional spatial mark-resight (SMR), generalized spatial mark-resight (gSMR), spatial capture-recapture (SCR), and hybrid SCR+gSMR models incorporating behavioral status (transient vs resident) in coyotes (*Canis latrans*).

In a post-hoc analysis, we allowed baseline detection rates and σ to vary as a function of behavioral status for the SMR+GPS, gSMR+GPS, SCR+GPS, and hybrid gSMR+SCR+GPS models for coyotes described in the main text. Our approach was analogous to the methods outlined by previous studies allowing these parameters to vary as a function of sex (e.g. Sollmann et al. 2011), except we replace sex with behavioral status. Behavioral status was only partially observed; that is, we could only classify the status of the marked segment of the population monitored with GPS collars. Animals were classified as residents if they were faithful to small (<20 km^2^) home ranges, and animals were classified as transients if they did not adhere to a home range and instead exhibited random, transitory movements or if calculated home range sizes were large (200–400 km^2^). To aid in the identifiability of parameter estimating transient or residency status (psi.trans), we assigned an informative prior distribution to the probability of being a transient as psi.trans~ beta(2.4, 7.2) which corresponds to *x̅* = 0.253 and SD = 0.133 probability of being a transient. These values were obtained by taking the mean and standard deviation of estimates of the proportion of transient coyotes in a given population from 11 previous studies (Bekoff 1978, Andelt 1985, Gese et al. 1988, Windberg and Knowlton 1988, Gese et al. 1999, Mills and Knowlton 1991, Windberg et al. 1997, Chamberlain et al. 2000, Berger and Gese 2007, Hinton et al. 2015, and Mitchell et al. 2015). We used moment-matching to find the parameters of the beta distribution matching this mean and SD using: $\alpha= \frac{(\mu^{2} -\mu^{3}-\sigma\mu^{2})}{\sigma^{2}}$ and $\beta= \frac{\mu-2\mu^{2}+\mu^{3}-\sigma^{2}+{\mu\sigma}^{2}}{\sigma^{2}}$ (Hobbs and Hooten 2015).
