## Supplementary material for "Evaluating and integrating spatial capture-recapture models with data of variable individual identifiability": sex model updates

**Appendix S6**: Tables including measures of central tendency, detection parameters, credible intervals, and coefficients of variation for sex-specific density estimation models for black bears, bobcats, cougars, and coyotes. Models include spatial count (SC), spatial mark resight (SMR), generalized spatial mark resight (gSMR), spatial capture recapture (SCR), and a novel hybrid model combining SCR and gSMR. Each model is fit with and without global positioning system (GPS) collar data from a subset of individuals within each population. See Appendix S4 for similar tables for base models, Appendix S7 for models with reduced sampling periods, and Appendix S8 for models incorporating behavioral status of coyotes.

**Table S1**: Sex-specific density estimates for black bears across a suite of models. Densities are presented as the number of animals per 100 km^2^. HPDI = highest posterior density interval. BCI = Bayesian Credible Interval. CV = coefficient of variation (defined as standard deviation divided by the posterior mean).

| **Model** | **Mean** | **Median** | **Mode** | **SD** | **Lower 95% HPDI** | **Upper 95% HPDI** | **Lower 95% BCI** | **Upper 95% BCI** | **CV** |
| --- | --- | --- | --- | --- | --- | --- | --- | --- | --- |
| SMR | 17.94 | 17.72 | 17.3 | 4.27 | 10.77 | 26.65 | 9.95 | 26.03 | 0.24 |
| SMR + GPS | 12.72 | 12.17 | 11.48 | 3.46 | 6.83 | 19.9 | 7.3 | 21.07 | 0.27 |
| gSMR | 21.32 | 21.09 | 20.72 | 5.33 | 11.15 | 31.22 | 11.88 | 32.55 | 0.25 |
| gSMR + GPS | 11.8 | 11.45 | 11.03 | 3.18 | 6.13 | 18.14 | 6.75 | 19.16 | 0.27 |
| SCR | 13.95 | 14.25 | 15.71 | 5.91 | 3.54 | 23.99 | 3.43 | 23.91 | 0.42 |
| SCR + GPS | 11.07 | 11.17 | 11.63 | 2.41 | 6.89 | 15.33 | 6.43 | 15.09 | 0.22 |
| SCR + gSMR | 19.72 | 19.59 | 18.81 | 4.33 | 11.28 | 27.76 | 11.43 | 27.96 | 0.22 |
| SCR + gSMR + GPS | 9.22 | 9.05 | 8.93 | 1.79 | 6.04 | 12.77 | 6.28 | 13.31 | 0.19 |

**Table S2**: Detection parameters for sex-specific models estimating the density of black bears. Values are presented as the median of the posterior distribution with 95% Bayesian Credible Intervals in parentheses. σ = spatial scale parameter, ${\lambda0}_{resight}$ = baseline detection rate for camera analyses, ${\lambda0}_{marking}$ = baseline detection rate for the marking process, $p0$ (intercept) = baseline detection probability on the logit scale for genetic SCR models when all covariates are at zero, $p0$ (survey effort) = effect of survey effort (distance traveled by scat detection dogs) on baseline detection probability in genetic SCR models on the logit scale, and $p0$(sex) = effect of being male (compared to females) on baseline detection probability in genetic SCR models on the logit scale.

| **Model** | **σ female** | **σ male** | ${\boldsymbol{\lambda}\boldsymbol{0}}_{\boldsymbol{resight}}$ **female** | ${\boldsymbol{\lambda}\boldsymbol{0}}_{\boldsymbol{resight}}$ **male** | ${\boldsymbol{\lambda}\boldsymbol{0}}_{\boldsymbol{marking}}$ | $\boldsymbol{p}\boldsymbol{0}$ **(intercept)** | $\boldsymbol{p}\boldsymbol{0}$ **(survey effort)** | $\boldsymbol{p}\boldsymbol{0}$ **(sex)** |
| --- | --- | --- | --- | --- | --- | --- | --- | --- |
| SMR | 560.89 (469.37–691.29) | 5269.79 (3289.43–12821.16) | 0.41 (0.24–0.62) | 0.02 (0.01–0.06) |  |  |  |  |
| SMR + GPS | 1953.52 (1828.38–2096.05) | 4084.05 (3863.52–4324.39) | 0.06 (0.03–0.12) | 0.02 (0.01–0.03) |  |  |  |  |
| gSMR | 574.28 (481.44–696.47) | 5550.45 (3340.04–9767.54) | 0.36 (0.22–0.54) | 0.03 (0.01–0.09) | 0.27 (0.09–0.77) |  |  |  |
| gSMR + GPS | 1956.63 (1829.93–2097.87) | 4077.95 (3864.93–4323.61) | 0.07 (0.03–0.13) | 0.02 (0.01–0.03) | 0.09 (0.03–0.2) |  |  |  |
| SCR | 1850.09 (706.43–18278.82) | 7461.1 (3786.02–18730.44) |  |  |  | -3.65 (-6.64–-1.42) | 0.43 (0.16–0.68) | -1.71 (-4.3–1.41) |
| SCR + GPS | 4475.59 (4190.24–4795) | 7537.76 (7130.07–7981.75) |  |  |  | -4.81 (-5.56–-4.04) | 0.42 (0.17–0.65) | -0.5 (-1.56–0.51) |
| gSMR + SCR | 827.36 (618.04–1261.23) | 5386.28 (3679.99–8982) | 0.14 (0.05–0.27) | 0.01 (0.01–0.02) | 0.07 (0.03–0.16) | -2.28 (-3.26–-1.38) | 0.44 (0.15–0.71) | -2.04 (-3.15–-0.96) |
| gSMR + SCR + GPS | 3647.2 (3406.83–3914.74) | 6136.59 (5813.99–6494.81) | 0.02 (0.01–0.04) | 0.01 (0.01–0.02) | 0.04 (0.02–0.08) | -4.24 (-5–-3.58) | 0.41 (0.14–0.64) | -0.36 (-1.24–0.55) |

**Table S3**: Sex-specific density estimates for bobcats across a suite of models. Densities are presented as the number of animals per 100 km^2^. HPDI = highest posterior density interval. BCI = Bayesian Credible Interval. CV = coefficient of variation (defined as the standard deviation divided by the posterior mean).

| **Model** | **Mean** | **Median** | **Mode** | **SD** | **Lower 95% HPDI** | **Upper 95% HPDI** | **Lower 95% BCI** | **Upper 95% BCI** | **CV** |
| --- | --- | --- | --- | --- | --- | --- | --- | --- | --- |
| SMR | 10.77 | 9.83 | 7.49 | 6.32 | 0.57 | 22.6 | 1.23 | 23.51 | 0.59 |
| SMR + GPS | 11.53 | 10.73 | 8.47 | 5.31 | 3.19 | 22.44 | 3.52 | 23.01 | 0.46 |
| gSMR | 10.16 | 9.75 | 8.92 | 5.72 | 0.33 | 20.39 | 0.82 | 22.03 | 0.56 |
| gSMR + GPS | 10.43 | 9.58 | 8.61 | 4.5 | 2.87 | 19.66 | 3.77 | 21.13 | 0.43 |
| SCR | 13.73 | 13.43 | 12.94 | 2.85 | 8.35 | 19.33 | 9.09 | 20.31 | 0.21 |
| SCR + GPS | 12.96 | 12.78 | 12.38 | 2.39 | 8.85 | 17.85 | 9.01 | 18.18 | 0.18 |
| SCR + gSMR | 12.61 | 12.53 | 12.27 | 2.07 | 8.85 | 16.71 | 8.93 | 16.95 | 0.16 |
| SCR + gSMR + GPS | 11.59 | 11.47 | 11.23 | 1.88 | 8.27 | 15.32 | 8.35 | 15.48 | 0.16 |

**Table S4**: Detection parameters for sex-specific models estimating the densities of bobcats. Values are presented as the median of the posterior distribution with 95% Bayesian Credible Intervals in parentheses. σ = spatial scale parameter, ${\lambda0}_{resight}$ = baseline detection rate for camera analyses, ${\lambda0}_{marking}$ = baseline detection rate for the marking process, $p0$ (intercept) = baseline detection probability on the logit scale for genetic SCR models when all covariates are at zero, $p0$ (survey effort) = effect of survey effort (distance traveled by scat detection dogs) on baseline detection probability in genetic SCR models on the logit scale, and $p0$(sex) = effect of being male (compared to females) on baseline detection probability in genetic SCR models on the logit scale.

| **Model** | **σ female** | **σ male** | ${\boldsymbol{\lambda}\boldsymbol{0}}_{\boldsymbol{resight}}$ **female** | ${\boldsymbol{\lambda}\boldsymbol{0}}_{\boldsymbol{resight}}$ **male** | ${\boldsymbol{\lambda}\boldsymbol{0}}_{\boldsymbol{marking}}$ | $\boldsymbol{p}\boldsymbol{0}$ **(intercept)** | $\boldsymbol{p}\boldsymbol{0}$ **(survey effort)** | $\boldsymbol{p}\boldsymbol{0}$ **(sex)** |
| --- | --- | --- | --- | --- | --- | --- | --- | --- |
| SMR | 2166.68 (663.6–15779.53) | 1048.94 (681.65–4895.78) | 0.01 (0–0.08) | 0.16 (0.04–2.32) |  |  |  |  |
| SMR + GPS | 1505.73 (1406.7–1617.32) | 2337.06 (2127.77–2584.98) | 0.01 (0–0.04) | 0.04 (0.02–0.08) |  |  |  |  |
| gSMR | 982.6 (578.57–5263.31) | 1488.66 (874.65–5555.84) | 0.04 (0.01–0.25) | 0.11 (0.04–1.12) | 0.45 (0.06–3.11) |  |  |  |
| gSMR + GPS | 1501.82 (1403.74–1606.86) | 2329.8 (2118.72–2579.92) | 0.02 (0–0.04) | 0.04 (0.02–0.09) | 0.11 (0.03–0.3) |  |  |  |
| SCR | 1485.62 (1213.18–1913.99) | 2065.89 (1462.97–3516.24) |  |  |  | -1.24 (-1.93–-0.55) | 0.46 (0.19–0.74) | -1.77 (-2.97–-0.65) |
| SCR + GPS | 1507.05 (1411.9–1615.81) | 2356.35 (2147.23–2596.73) |  |  |  | -1.26 (-1.81–-0.76) | 0.46 (0.21–0.73) | -1.96 (-2.8–-1.16) |
| gSMR + SCR | 1510.3 (1240.06–1937.53) | 1650.86 (1305.83–2232.81) | 0.01 (0–0.03) | 0.03 (0.02–0.05) | 0.13 (0.04–0.38) | -1.38 (-2.03–-0.75) | 0.49 (0.22–0.8) | -0.98 (-1.9–-0.05) |
| gSMR + SCR + GPS | 1505.35 (1415.79–1608.89) | 2311.92 (2117.05–2533.6) | 0.01 (0–0.03) | 0.02 (0.01–0.04) | 0.08 (0.02–0.21) | -1.37 (-1.89–-0.86) | 0.47 (0.22–0.75) | -1.52 (-2.27–-0.79) |

**Table S5**: Sex-specific density estimates for cougars across a suite of models. Densities are presented as the number of animals per 100 km^2^. HPDI = highest posterior density interval. BCI = Bayesian Credible Interval. CV = coefficient of variation (defined as the standard deviation divided by the posterior mean).

| **Model** | **Mean** | **Median** | **Mode** | **SD** | **Lower 95% HPDI** | **Upper 95% HPDI** | **Lower 95% BCI** | **Upper 95% BCI** | **CV** |
| --- | --- | --- | --- | --- | --- | --- | --- | --- | --- |
| SMR | 1.05 | 0.7 | 0.2 | 0.97 | 0.12 | 3.08 | 0.16 | 3.59 | 0.92 |
| SMR + GPS | 2.25 | 2.15 | 1.91 | 0.83 | 0.82 | 3.86 | 0.94 | 4.1 | 0.37 |
| gSMR | 2.32 | 2.19 | 1.86 | 0.92 | 0.82 | 4.18 | 0.94 | 4.53 | 0.4 |
| gSMR + GPS | 2.19 | 2.07 | 1.95 | 0.76 | 0.94 | 3.79 | 1.01 | 3.98 | 0.35 |
| SCR | 1.48 | 1.32 | 0.97 | 0.83 | 0.23 | 3.08 | 0.4 | 3.61 | 0.56 |
| SCR + GPS | 2.51 | 2.25 | 1.78 | 1.18 | 0.76 | 4.97 | 0.89 | 5.33 | 0.47 |
| SCR + gSMR | 2.52 | 2.42 | 2.19 | 0.87 | 0.96 | 4.2 | 1.19 | 4.6 | 0.35 |
| SCR + gSMR + GPS | 2.39 | 2.28 | 2.16 | 0.69 | 1.22 | 3.81 | 1.29 | 3.97 | 0.29 |

**Table S6**: Detection parameters for sex-specific models estimating the densities of cougars. Values are presented as the median of the posterior distribution with 95% Bayesian Credible Intervals in parentheses. σ = spatial scale parameter, ${\lambda0}_{resight}$ = baseline detection rate for camera analyses, ${\lambda0}_{marking}$ = baseline detection rate for the marking process, $p0$ (intercept) = baseline detection probability on the logit scale for genetic SCR models when all covariates are at zero, $p0$ (survey effort) = effect of survey effort (distance traveled by scat detection dogs) on baseline detection probability in genetic SCR models on the logit scale, and $p0$(sex) = effect of being male (compared to females) on baseline detection probability in genetic SCR models on the logit scale.

| **Model** | **σ female** | **σ male** | ${\boldsymbol{\lambda}\boldsymbol{0}}_{\boldsymbol{resight}}$ **female** | ${\boldsymbol{\lambda}\boldsymbol{0}}_{\boldsymbol{resight}}$ **male** | ${\boldsymbol{\lambda}\boldsymbol{0}}_{\boldsymbol{marking}}$ | $\boldsymbol{p}\boldsymbol{0}$ **(intercept)** | $\boldsymbol{p}\boldsymbol{0}$ **(survey effort)** | $\boldsymbol{p}\boldsymbol{0}$ **(sex)** |
| --- | --- | --- | --- | --- | --- | --- | --- | --- |
| SMR | 6388.18 (2530.12–15281.61) | 5377.48 (2182.98–18565.03) | 0.07 (0.01–3.6) | 0.04 (0.01–0.26) |  |  |  |  |
| SMR + GPS | 3135.1 (2932.13–3365.81) | 4799.03 (4533.41–5087.11) | 0.03 (0.02–0.06) | 0.02 (0.01–0.04) |  |  |  |  |
| gSMR | 2901.96 (2007.23–6257.69) | 4407.54 (2442.08–9075.38) | 0.04 (0.02–0.1) | 0.03 (0.01–0.08) | 0.48 (0.17–1.28) |  |  |  |
| gSMR + GPS | 3250.23 (3042.22–3482.17) | 4891.93 (4635.82–5192.51) | 0.03 (0.02–0.06) | 0.02 (0.01–0.04) | 0.43 (0.16–1.07) |  |  |  |
| SCR | 1175 (667.18–15266.91) | 5164.32 (2622.39–8976.27) |  |  |  | -4.82 (-10.28–-0.27) | 10.87 (0.45–24.71) | 6.72 (0.12–15.93) |
| SCR + GPS | 3152.54 (2947.77–3390.86) | 4811.07 (4556.17–5080.18) |  |  |  | -4.22 (-5.98–-2.76) | 0.57 (0.08–1.06) | 0.81 (-1.02–2.66) |
| gSMR + SCR | 2835.95 (2090.65–5259.9) | 4596.02 (3020.71–7868.03) | 0.04 (0.02–0.08) | 0.03 (0.01–0.09) | 0.44 (0.16–1.1) | -4.21 (-5.79–-2.96) | 0.63 (0.1–1.3) | 1.36 (-0.45–3.55) |
| gSMR + SCR + GPS | 3251.37 (3045.38–3489.19) | 4889.69 (4621.46–5174.91) | 0.03 (0.02–0.06) | 0.02 (0.01–0.04) | 0.41 (0.16–0.95) | -4.32 (-5.72–-3.18) | 0.58 (0.06–1.1) | 1.21 (-0.38–2.92) |

**Table S7**: Sex-specific density estimates for coyotes across a suite of models. Densities are presented as the number of animals per 100 km^2^. HPDI = highest posterior density interval. BCI=Bayesian Credible Interval. CV=coefficient of variation (defined as the standard deviation divided by the posterior mean).

| **Model** | **Mean** | **Median** | **Mode** | **SD** | **Lower 95% HPDI** | **Upper 95% HPDI** | **Lower 95% BCI** | **Upper 95% BCI** | **CV** |
| --- | --- | --- | --- | --- | --- | --- | --- | --- | --- |
| SMR | 48.68 | 48.22 | 47.94 | 7.73 | 34.18 | 64.24 | 34.79 | 65.15 | 0.16 |
| SMR + GPS | 34.31 | 33.87 | 33.5 | 5.56 | 24.11 | 45.77 | 24.87 | 47 | 0.16 |
| gSMR | 43.01 | 42.72 | 42.44 | 6.7 | 30.67 | 56.91 | 30.97 | 57.37 | 0.16 |
| gSMR + GPS | 31.24 | 30.82 | 30.56 | 4.65 | 21.97 | 39.98 | 23.04 | 41.5 | 0.15 |
| SCR | 33.51 | 33.24 | 32.93 | 3.46 | 27.31 | 40.55 | 27.44 | 40.83 | 0.1 |
| SCR + GPS | 24.67 | 24.55 | 24.33 | 2.42 | 20.41 | 29.66 | 20.41 | 29.66 | 0.1 |
| SCR + gSMR | 37.58 | 37.38 | 37.09 | 3.65 | 30.07 | 44.55 | 30.76 | 45.24 | 0.1 |
| SCR + gSMR + GPS | 28.12 | 28 | 27.79 | 2.54 | 23.59 | 33.38 | 23.45 | 33.38 | 0.09 |

**Table S8**: Detection parameters for sex-specific models estimating the densities of coyotes. Values are presented as the median of the posterior distribution with 95% Bayesian Credible Intervals in parentheses. σ = spatial scale parameter, ${\lambda0}_{resight}$ = baseline detection rate for camera analyses, ${\lambda0}_{marking}$ = baseline detection rate for the marking process, $p0$ (intercept) = baseline detection probability on the logit scale for genetic SCR models when all covariates are at zero, $p0$ (survey effort) = effect of survey effort (distance traveled by scat detection dogs) on baseline detection probability in genetic SCR models on the logit scale, and $p0$(sex) = effect of being male (compared to females) on baseline detection probability in genetic SCR models on the logit scale.

| **Model** | **σ female** | **σ male** | ${\boldsymbol{\lambda}\boldsymbol{0}}_{\boldsymbol{resight}}$ **female** | ${\boldsymbol{\lambda}\boldsymbol{0}}_{\boldsymbol{resight}}$ **male** | ${\boldsymbol{\lambda}\boldsymbol{0}}_{\boldsymbol{marking}}$ | $\boldsymbol{p}\boldsymbol{0}$ **(intercept)** | $\boldsymbol{p}\boldsymbol{0}$ **(survey effort)** | $\boldsymbol{p}\boldsymbol{0}$ **(sex)** |
| --- | --- | --- | --- | --- | --- | --- | --- | --- |
| SMR | 439.7 (385.31–497.28) | 937.69 (792.48–1134.37) | 1.07 (0.78–1.41) | 0.22 (0.14–0.35) |  |  |  |  |
| SMR + GPS | 3225.55 (3011.37–3460.18) | 2436.67 (2351.52–2526.31) | 0.02 (0.01–0.04) | 0.06 (0.04–0.08) |  |  |  |  |
| gSMR | 516.67 (465.44–582.56) | 1090.28 (943.15–1292.26) | 0.9 (0.66–1.19) | 0.18 (0.12–0.27) | 0.18 (0.08–0.36) |  |  |  |
| gSMR + GPS | 3220.71 (3021.79–3435.5) | 2460.63 (2371.67–2549.16) | 0.02 (0.01–0.04) | 0.06 (0.05–0.08) | 0.02 (0.01–0.04) |  |  |  |
| SCR | 1175.68 (1006.23–1418.81) | 1381.25 (1184.39–1629.71) |  |  |  | -1.24 (-1.82–-0.67) | 0.61 (0.37–0.94) | -0.35 (-1.06–0.39) |
| SCR + GPS | 3130.83 (2933.49–3344.39) | 2433.62 (2347.76–2525.48) |  |  |  | -2.96 (-3.39–-2.56) | 0.4 (0.24–0.55) | 0.39 (-0.13–0.92) |
| gSMR + SCR | 1016.22 (841.6–1240.43) | 1258.42 (1103.38–1427.17) | 0.23 (0.06–0.42) | 0.16 (0.09–0.29) | 0.07 (0.04–0.14) | -1.06 (-1.64–-0.46) | 0.62 (0.37–0.94) | -0.33 (-1.06–0.31) |
| gSMR + SCR + GPS | 3069.76 (2871.63–3277.89) | 2422.85 (2340.07–2507.23) | 0.03 (0.01–0.05) | 0.07 (0.05–0.09) | 0.02 (0.01–0.03) | -3 (-3.37–-2.63) | 0.39 (0.23–0.53) | 0.29 (-0.19–0.79) |
