## Supplementary material for "Evaluating and integrating spatial capture-recapture models with data of variable individual identifiability": reduced sampling model tables

**Appendix S7**: Tables including measures of central tendency, detection parameters, credible intervals, and coefficients of variation for reduced sampling period density estimation models for black bears, bobcats, cougars, and coyotes. The results presented here are for analyses using cameras with a sampling period reduced from 140 days to 70 days. Models include spatial count (SC), spatial mark-resight (SMR), generalized spatial mark-resight (gSMR), spatial capture-recapture (SCR), and a novel hybrid model combining SCR and gSMR. Each model is fit with and without global positioning system (GPS) collar data from a subset of individuals within each population. See Appendix S4 for similar tables for base models, Appendix S6 for sex-specific models, and Appendix S8 for models incorporating behavioral status of coyotes.

**Table S1**: Density estimates for reduced sampling period models for black bears across a suite of models. Densities are presented as the number of animals per 100 km^2^. HPDI = highest posterior density interval. BCI = Bayesian Credible Interval. CV = coefficient of variation (defined as the standard deviation divided by the posterior mean).

| **Model** | **Mean** | **Median** | **Mode** | **SD** | **Lower 95% HPDI** | **Upper 95% HPDI** | **Lower 95% BCI** | **Upper 95% BCI** | **CV** |
| --- | --- | --- | --- | --- | --- | --- | --- | --- | --- |
| SC | 211.97 | 176.22 | 88.95 | 133.19 | 33.72 | 473.19 | 48.95 | 506.92 | 0.63 |
| SC + GPS | 1.46 | 0.74 | 0.39 | 2.14 | 0.04 | 5.58 | 0.2 | 9.41 | 1.47 |
| SPA | 219.33 | 189.28 | 80.12 | 142.89 | 17.95 | 482.44 | 32.09 | 505.83 | 0.65 |
| SPA + GPS | 1.42 | 0.86 | 0.47 | 1.49 | 0.08 | 5.07 | 0.2 | 6.24 | 1.05 |
| SMR | 7.55 | 7.53 | 7.39 | 3.03 | 2.65 | 13.5 | 2.15 | 13.15 | 0.4 |
| SMR + GPS | 14.08 | 13.66 | 13.22 | 3.65 | 7.1 | 21.66 | 8.19 | 23.02 | 0.26 |
| gSMR | 8.37 | 8.39 | 8.44 | 2.85 | 3.39 | 13.62 | 2.73 | 13.27 | 0.34 |
| gSMR + GPS | 11.81 | 11.51 | 10.99 | 2.82 | 6.63 | 17.33 | 7.1 | 17.99 | 0.24 |
| SCR + gSMR | 12.25 | 12.13 | 11.93 | 2.85 | 7.12 | 18.14 | 7.01 | 18.06 | 0.23 |
| SCR + gSMR + GPS | 8.94 | 8.82 | 8.57 | 1.72 | 5.89 | 12.51 | 6.08 | 12.82 | 0.19 |

**Table S2**: Detection parameters for density models with a reduced sampling period for black bears. Values are presented as the median of the posterior distribution with 95% Bayesian Credible Intervals in parentheses. σ = spatial scale parameter, ${\lambda0}_{resight}$ = baseline detection rate for camera analyses, ${\lambda0}_{marking}$ = baseline detection rate for the marking process, $p0$ (intercept) = baseline detection probability on the logit scale for genetic SCR models when all covariates are at zero, $p0$ (survey effort) = effect of survey effort (distance traveled by scat detection dogs) on baseline detection rate in genetic SCR models on the logit scale.

| **Model** | **σ** | ${\boldsymbol{\lambda}\boldsymbol{0}}_{\boldsymbol{resight}}$ | ${\boldsymbol{\lambda}\boldsymbol{0}}_{\boldsymbol{marking}}$ | $\boldsymbol{p}\boldsymbol{0}$ **(intercept)** | $\boldsymbol{p}\boldsymbol{0}$ **(survey effort)** |
| --- | --- | --- | --- | --- | --- |
| SC | 199.51 (105.95–377.74) | 0.69 (0.41–1.18) |  |  |  |
| SC + GPS | 6486.45 (6218.57–6777.43) | 0.21 (0.01–1.44) |  |  |  |
| SPA | 288.86 (140.9-826.71) | 0.23 (0.06–0.44) |  |  |  |
| SPA + GPS | 6490.86 (6219.73–6785.74) | 0.12 (0.01–0.67) |  |  |  |
| SMR | 4515.35 (3082.51–8400.48) | 0.03 (0.02–0.07) |  |  |  |
| SMR + GPS | 3393.82 (3247.93–3543.98) | 0.03 (0.02–0.05) |  |  |  |
| gSMR | 3826.74 (2776.88–6261.02) | 0.04 (0.02–0.08) | 0.07 (0.03–0.19) |  |  |
| gSMR + GPS | 3394.4 (3249.29–3546.78) | 0.04 (0.02–0.05) | 0.07 (0.03–0.14) |  |  |
| gSMR + SCR | 3955.49 (3050.38–5531.83) | 0.02 (0.02–0.04) | 0.04 (0.02–0.09) | -4.34 (-4.9–-3.84) | 0.41 (0.15–0.66) |
| gSMR + SCR + GPS | 5283.01 (5060.83–5519.62) | 0.02 (0.01–0.03) | 0.03 (0.01–0.07) | -4.62 (-5.1–-4.18) | 0.41 (0.14–0.65) |

**Table S3:** Density estimates for reduced sampling period models for bobcats across a suite of models. Densities are presented as the number of animals per 100 km^2^. HPDI = highest posterior density interval. BCI = Bayesian Credible Interval. CV = coefficient of variation (defined as the standard deviation divided by the posterior mean).

| **Model** | **Mean** | **Median** | **Mode** | **SD** | **Lower 95% HPDI** | **Upper 95% HPDI** | **Lower 95% BCI** | **Upper 95% BCI** | **CV** |
| --- | --- | --- | --- | --- | --- | --- | --- | --- | --- |
| SC | 93.35 | 84.14 | 24.51 | 69.5 | 0.47 | 217.77 | 0.94 | 226.74 | 0.74 |
| SC + GPS | 49.13 | 36.29 | 10.79 | 39.62 | 1.89 | 126.8 | 3.3 | 133.4 | 0.81 |
| SPA | 97.77 | 91.8 | 41.68 | 68.51 | 0.29 | 214.58 | 0.58 | 222.4 | 0.7 |
| SPA + GPS | 48 | 31.11 | 9.66 | 44.47 | 1.41 | 146.12 | 3.3 | 156.49 | 0.93 |
| SMR | 7.16 | 6.22 | 4.82 | 4.68 | 0.33 | 16.22 | 0.82 | 18.35 | 0.65 |
| SMR + GPS | 8.11 | 7.37 | 6.49 | 4.03 | 1.97 | 16.22 | 2.62 | 18.84 | 0.5 |
| gSMR | 5.71 | 5.16 | 4.36 | 3.66 | 0.49 | 12.69 | 0.82 | 14.74 | 0.64 |
| gSMR + GPS | 7.46 | 6.88 | 6.12 | 3.08 | 2.29 | 13.43 | 2.95 | 14.91 | 0.41 |
| SCR + gSMR | 12.43 | 12.29 | 12.18 | 2.04 | 8.52 | 16.46 | 8.85 | 16.95 | 0.16 |
| SCR + gSMR + GPS | 12.06 | 11.88 | 11.6 | 2.03 | 8.19 | 15.97 | 8.52 | 16.38 | 0.17 |

**Table S4**: Detection parameters for density models with a reduced sampling period for bobcats. Values are presented as the median of the posterior distribution with 95% Bayesian Credible Intervals in parentheses. σ = spatial scale parameter, ${\lambda0}_{resight}$ = baseline detection rate for camera analyses, ${\lambda0}_{marking}$ = baseline detection rate for the marking process, $p0$ (intercept) = baseline detection probability on the logit scale for genetic SCR models when all covariates are at zero, $p0$ (survey effort) = effect of survey effort (distance traveled by scat detection dogs) on baseline detection rate in genetic SCR models on the logit scale.

| **Model** | **σ** | ${\boldsymbol{\lambda}\boldsymbol{0}}_{\boldsymbol{resight}}$ | ${\boldsymbol{\lambda}\boldsymbol{0}}_{\boldsymbol{marking}}$ | $\boldsymbol{p}\boldsymbol{0}$ **(intercept)** | $\boldsymbol{p}\boldsymbol{0}$ **(survey effort)** |
| --- | --- | --- | --- | --- | --- |
| SC | 475.33 (106.76–17124.04) | 0.03 (0–0.31) |  |  |  |
| SC + GPS | 1829.84 (1730.92–1941.85) | 0 (0–0.04) |  |  |  |
| SPA | 422.74 (127.9–17498.85) | 0.04 (0–0.31) |  |  |  |
| SPA + GPS | 1829.34 (1726.29–1935.73) | 0.01 (0–0.07) |  |  |  |
| SMR | 1786.87 (966.1–6181.45) | 0.04 (0.01–0.1) |  |  |  |
| SMR + GPS | 1833.82 (1735.24–1940.59) | 0.03 (0.01–0.06) |  |  |  |
| gSMR | 1871.11 (1066.05–4825.41) | 0.04 (0.01–0.14) | 0.23 (0.04–1.25) |  |  |
| gSMR + GPS | 1826.77 (1734.3–1933.84) | 0.03 (0.01–0.06) | 0.12 (0.03–0.31) |  |  |
| gSMR + SCR | 1654.18 (1405.55–2005.92) | 0.02 (0.01–0.04) | 0.12 (0.03–0.31) | -1.98 (-2.45–-1.52) | 0.45 (0.2–0.75) |
| gSMR + SCR + GPS | 1825.5 (1732.81–1932.5) | 0.02 (0.01–0.03) | 0.08 (0.02–0.2) | -2.16 (-2.55–-1.78) | 0.45 (0.2–0.73) |

**Table S5**: Density estimates for reduced sampling period models for cougars across a suite of models. Densities are presented as the number of animals per 100 km^2^. HPDI = highest posterior density interval. BCI = Bayesian Credible Interval. CV = coefficient of variation (defined as the standard deviation divided by the posterior mean).

| **Model** | **Mean** | **Median** | **Mode** | **SD** | **Lower 95% HPDI** | **Upper 95% HPDI** | **Lower 95% BCI** | **Upper 95% BCI** | **CV** |
| --- | --- | --- | --- | --- | --- | --- | --- | --- | --- |
| SC | 156.94 | 141.23 | 92.3 | 100.39 | 1.46 | 336.68 | 9.1 | 351.24 | 0.64 |
| SC + GPS | 5.44 | 3.82 | 0.93 | 4.85 | 0.04 | 15.8 | 0.35 | 17.52 | 0.89 |
| SPA | 25.31 | 16.78 | 1.81 | 24.05 | 0.08 | 71.87 | 0.25 | 75.96 | 0.95 |
| SPA + GPS | 5.63 | 3.98 | 0.92 | 4.89 | 0.12 | 15.8 | 0.35 | 17.48 | 0.87 |
| SMR | 2 | 1.76 | 0.25 | 1.54 | 0.12 | 4.96 | 0.16 | 5.7 | 0.77 |
| SMR + GPS | 2.02 | 1.87 | 1.72 | 0.83 | 0.7 | 3.63 | 0.78 | 3.98 | 0.41 |
| gSMR | 2.44 | 2.26 | 1.94 | 1.08 | 0.55 | 4.45 | 0.9 | 5.07 | 0.44 |
| gSMR + GPS | 1.7 | 1.6 | 1.49 | 0.6 | 0.66 | 2.89 | 0.78 | 3.12 | 0.35 |
| SCR + gSMR | 2.35 | 2.22 | 1.97 | 0.8 | 0.99 | 3.94 | 1.13 | 4.17 | 0.34 |
| SCR + gSMR + GPS | 1.88 | 1.82 | 1.65 | 0.54 | 0.93 | 2.95 | 1.03 | 3.08 | 0.29 |

**Table S6**: Detection parameters for density models with a reduced sampling period for cougars. Values are presented as the median of the posterior distribution with 95% Bayesian Credible Intervals in parentheses. σ = spatial scale parameter, ${\lambda0}_{resight}$ = baseline detection rate for camera analyses, ${\lambda0}_{marking}$ = baseline detection rate for the marking process, $p0$ (intercept) = baseline detection probability on the logit scale for genetic SCR models when all covariates are at zero, $p0$ (survey effort) = effect of survey effort (distance traveled by scat detection dogs) on baseline detection rate in genetic SCR models on the logit scale.

| **Model** | **σ** | ${\boldsymbol{\lambda}\boldsymbol{0}}_{\boldsymbol{resight}}$ | ${\boldsymbol{\lambda}\boldsymbol{0}}_{\boldsymbol{marking}}$ | $\boldsymbol{p}\boldsymbol{0}$ **(intercept)** | $\boldsymbol{p}\boldsymbol{0}$ **(survey effort)** |
| --- | --- | --- | --- | --- | --- |
| SC | 254.91 (106.39–1671.55) | 0.1 (0.01–0.36) |  |  |  |
| SC + GPS | 5068.05 (4877.84–5282.71) | 0.01 (0–0.11) |  |  |  |
| SPA | 2818.17 (349.01–18561.7) | 0.01 (0–0.12) |  |  |  |
| SPA + GPS | 5066.44 (4867.55–5268.41) | 0.01 (0–0.09) |  |  |  |
| SMR | 4365.04 (2216.4–18414.61) | 0.03 (0.01–0.07) |  |  |  |
| SMR + GPS | 5055.87 (4862.87–5255.02) | 0.02 (0.01–0.03) |  |  |  |
| gSMR | 3547.05 (2392.2–6451.37) | 0.03 (0.01–0.06) | 0.39 (0.13–1.19) |  |  |
| gSMR + GPS | 5196.2 (5002.76–5410.99) | 0.02 (0.01–0.03) | 0.32 (0.11–0.81) |  |  |
| gSMR + SCR | 3630.5 (2658.02–5739.51) | 0.03 (0.02–0.06) | 0.4 (0.15–1.06) | -3.53 (-4.44–-2.69) | 0.54 (0.04–1.04) |
| gSMR + SCR + GPS | 4432.37 (4258.04–4611.01) | 0.03 (0.02–0.04) | 0.43 (0.15–1.02) | -3.69 (-4.48–-3.02) | 0.53 (0.07–1.02) |

**Table S7**: Density estimates for reduced sampling period models for coyotes across a suite of models. Densities are presented as the number of animals per 100 km^2^. HPDI = highest posterior density interval. BCI = Bayesian Credible Interval. CV = coefficient of variation (defined as the standard deviation divided by the posterior mean).

| **Model** | **Mean** | **Median** | **Mode** | **SD** | **Lower 95% HPDI** | **Upper 95% HPDI** | **Lower 95% BCI** | **Upper 95% BCI** | **CV** |
| --- | --- | --- | --- | --- | --- | --- | --- | --- | --- |
| SC | 125.15 | 115.01 | 95.53 | 50.32 | 50.91 | 223.43 | 52.32 | 226.73 | 0.4 |
| SC + GPS | 3.42 | 2.75 | 1.86 | 2.49 | 0.61 | 7.32 | 1.07 | 9.61 | 0.73 |
| SPA | 108.44 | 97.1 | 82.67 | 49.95 | 33.47 | 214 | 38.65 | 222.02 | 0.46 |
| SPA + GPS | 4.7 | 3.77 | 2.37 | 2.92 | 1.89 | 10.37 | 1.89 | 12.26 | 0.62 |
| SMR | 35.85 | 35.32 | 34.82 | 6.72 | 22.12 | 48.22 | 24.11 | 50.66 | 0.19 |
| SMR + GPS | 22.69 | 22.28 | 21.77 | 4.38 | 14.5 | 31.28 | 15.11 | 32.35 | 0.19 |
| gSMR | 36.55 | 35.7 | 34.74 | 7.26 | 24.11 | 51.42 | 24.41 | 52.34 | 0.2 |
| gSMR + GPS | 25.48 | 25.02 | 24.5 | 4.47 | 17.24 | 34.48 | 17.85 | 35.4 | 0.18 |
| SCR | 33.96 | 33.79 | 33.47 | 3.26 | 27.86 | 40.28 | 28.28 | 40.69 | 0.1 |
| SCR + GPS | 25.51 | 25.38 | 25.2 | 2.24 | 21.24 | 29.93 | 21.38 | 30.34 | 0.09 |
| SCR + gSMR | 125.15 | 115.01 | 95.53 | 50.32 | 50.91 | 223.43 | 52.32 | 226.73 | 0.4 |
| SCR + gSMR + GPS | 3.42 | 2.75 | 1.86 | 2.49 | 0.61 | 7.32 | 1.07 | 9.61 | 0.73 |

**Table S8**: Detection parameters for density models with a reduced sampling period for coyotes. Values are presented as the median of the posterior distribution with 95% Bayesian Credible Intervals in parentheses. σ = spatial scale parameter, ${\lambda0}_{resight}$ = baseline detection rate for camera analyses, ${\lambda0}_{marking}$ = baseline detection rate for the marking process, $p0$ (intercept) = baseline detection probability on the logit scale for genetic SCR models when all covariates are at zero, $p0$ (survey effort) = effect of survey effort (distance traveled by scat detection dogs) on baseline detection rate in genetic SCR models on the logit scale.

| **Model** | **σ** | ${\boldsymbol{\lambda}\boldsymbol{0}}_{\boldsymbol{resight}}$ | ${\boldsymbol{\lambda}\boldsymbol{0}}_{\boldsymbol{marking}}$ | $\boldsymbol{p}\boldsymbol{0}$ **(intercept)** | $\boldsymbol{p}\boldsymbol{0}$ **(survey effort)** |
| --- | --- | --- | --- | --- | --- |
| SC | 278.19 (181.29–416.37) | 0.99 (0.67–1.44) |  |  |  |
| SC + GPS | 2671.04 (2585.92–2760.22) | 0.6 (0.16–1.18) |  |  |  |
| SPA | 384.45 (225.17–642.21) | 0.44 (0.24–0.66) |  |  |  |
| SPA + GPS | 2729.21 (2644.5–2818.39) | 0.29 (0.09–0.61) |  |  |  |
| SMR | 656.43 (578.58–759.43) | 0.62 (0.42–0.9) |  |  |  |
| SMR + GPS | 2639.39 (2560.32–2729.72) | 0.06 (0.04–0.08) |  |  |  |
| gSMR | 849 (730.45–1018.26) | 0.36 (0.24–0.53) | 0.15 (0.06–0.31) |  |  |
| gSMR + GPS | 2656.85 (2574.93–2742.68) | 0.05 (0.04–0.07) | 0.02 (0.01–0.04) |  |  |
| gSMR + SCR | 1272.99 (1158.08–1408.9) | 0.16 (0.12–0.2) | 0.07 (0.03–0.12) | -1.41 (-1.73–-1.1) | 0.6 (0.38–0.89) |
| gSMR + SCR + GPS | 2595.27 (2511.87–2677.82) | 0.05 (0.04–0.06) | 0.02 (0.01–0.04) | -2.71 (-2.94–-2.48) | 0.38 (0.23–0.53) |
