## Supplementary material for "Evaluating and integrating spatial capture-recapture models with data of variable individual identifiability": behavioral status model tables

**Appendix S8**: Tables including measures of central tendency, detection parameters, credible intervals, and coefficients of variation for density estimation models incorporating behavioral status for coyotes. See Appendix S5 for details on methods. Models include spatial count (SC), spatial mark-resight (SMR), generalized spatial mark-resight (gSMR), spatial capture-recapture (SCR), and a novel hybrid model combining SCR and gSMR. Each model is fit with and without global positioning system (GPS) collar data from a subset of individuals within each population. See Appendix S4 for similar tables for base models, Appendix S6 for sex-specific models, and Appendix S7 for models with reduced sampling periods.

**Table S1**: Behavioral status-specific density estimates for coyotes across a suite of models. Densities are presented as the number of animals per 100 km^2^. HPDI = highest posterior density interval. BCI = Bayesian Credible Interval. CV = coefficient of variation (defined as standard deviation divided by the posterior mean).

| **Model** | **Mean** | **Median** | **Mode** | **SD** | **Lower 95% HPDI** | **Upper 95% HPDI** | **Lower 95% BCI** | **Upper 95% BCI** | **CV** |
| --- | --- | --- | --- | --- | --- | --- | --- | --- | --- |
| SMR | 53.32 | 52.79 | 52.39 | 9.55 | 36.01 | 72.93 | 35.55 | 72.78 | 0.18 |
| SMR + GPS | 44.83 | 44.71 | 45.03 | 7.55 | 31.13 | 60.58 | 31.28 | 60.88 | 0.17 |
| gSMR | 42.88 | 42.11 | 40.87 | 7.19 | 30.21 | 57.83 | 30.67 | 58.29 | 0.17 |
| gSMR + GPS | 39.75 | 39.37 | 38.93 | 6.14 | 27.62 | 51.42 | 29.14 | 52.95 | 0.15 |
| SCR | 40.14 | 39.03 | 37.38 | 7.57 | 27.17 | 55.59 | 28.55 | 57.8 | 0.19 |
| SCR + GPS | 40.53 | 39.03 | 37.56 | 7.86 | 27.86 | 57.38 | 28.97 | 58.76 | 0.19 |
| SCR + gSMR | 40.35 | 39.72 | 38.48 | 5.64 | 30.48 | 51.59 | 31.03 | 52.41 | 0.14 |
| SCR + gSMR + GPS | 34.60 | 34.34 | 33.87 | 3.28 | 28.00 | 40.69 | 28.55 | 41.52 | 0.09 |

**Table S2**: Detection parameters for density models incorporating behavioral status in coyotes. Values are presented as the median of the posterior distribution with 95% Bayesian Credible Intervals in parentheses. σ = spatial scale parameter, ${\lambda0}_{resight}$ = baseline detection rate for camera analyses, ${\lambda0}_{marking}$ = baseline detection rate for the marking process, $p0$ (intercept) = baseline detection probability on the logit scale for genetic SCR models when all covariates are at zero, $p0$ (survey effort) = effect of survey effort (distance traveled by scat detection dogs) on baseline detection probability in genetic SCR models on the logit scale, and $p0$(behavioral status) = effect of being transient (compared to resident) on baseline detection probability in genetic SCR models on the logit scale.

| **Model** | **σ resident** | **σ transient** | ${\boldsymbol{\lambda}\boldsymbol{0}}_{\boldsymbol{resight}}$ **resident** | ${\boldsymbol{\lambda}\boldsymbol{0}}_{\boldsymbol{resight}}$ **transient** | ${\boldsymbol{\lambda}\boldsymbol{0}}_{\boldsymbol{marking}}$ | $\boldsymbol{p}\boldsymbol{0}$ **(intercept)** | $\boldsymbol{p}\boldsymbol{0}$ **(survey effort)** | $\boldsymbol{p}\boldsymbol{0}$ **(behavioral status)** |
| --- | --- | --- | --- | --- | --- | --- | --- | --- |
| SMR | 633.99 (580.66-699.24) | 5934.39 (544.08-18933.24) | 0.63 (0.48–0.8) | 0 (0–0.06) |  |  |  |  |
| SMR + GPS | 1058.92 (1020.35-1102.84) | 4384.62 (4140.7-4649.14) | 0.24 (0.18–0.3) | 0 (0–0.01) |  |  |  |  |
| gSMR | 696.73 (638.86-769.54) | 1706.86 (1093.21-3746.16) | 0.54 (0.4–0.7) | 0.02 (0–0.05) | 0.12 (0.05–0.26) |  |  |  |
| gSMR + GPS | 1059.48 (1020.24-1101.25) | 4389.48 (4160.52-4640.23) | 0.22 (0.18–0.28) | 0.01 (0–0.02) | 0.05 (0.02–0.1) |  |  |  |
| SCR | 1195.13 (1073.81-1353.4) | 8883.97 (2469.15-19294.17) |  |  |  | -0.93 (-1.34– -0.48) | 0.64 (0.4–0.94) | -5.53 (-6.97– -3.48) |
| SCR + GPS | 1102.62 (1063.24-1146.04) | 4371.06 (4139.84-4651.25) |  |  |  | -0.76 (-1.11– -0.39) | 0.7 (0.45–0.99) | -4.8 (-6.13– -3.74) |
| gSMR + SCR | 893.03 (741.08-1077.34) | 1738.18 (1310.21-3970.42) | 0.35 (0.23–0.54) | 0.01 (0–0.03) | 0.06 (0.03–0.13) | -0.96 (-1.71– -0.46) | 0.7 (0.42–1.1) | -0.76 (-4–0.63) |
| gSMR + SCR + GPS | 1077.43 (1039.95-1117.08) | 4394.1 (4151.32-4638.7) | 0.23 (0.19–0.28) | 0.01 (0–0.02) | 0.04 (0.02–0.09) | -0.98 (-1.27– -0.68) | 0.75 (0.48–1.1) | -3.44 (-4.58– -2.46) |
